# Pharmacological induction of mitochondrial stress counteracts therapy resistance in glioblastoma stem-like cells

**DOI:** 10.64898/2026.02.05.704021

**Authors:** Kenji Miki, Maged T. Ghoche, Fanen Yuan, Fengping Li, Namya Manoj, Skyler Oken, Koji Yoshimoto, Pascal O. Zinn, Sameer Agnihotri, Samuel K. McBrayer, Kalil G. Abdullah

**Affiliations:** Department of Neurosurgery, University of Pittsburgh School of Medicine, Pittsburgh, PA, USA; Hillman Comprehensive Cancer Center, University of Pittsburgh Medical Center, Pittsburgh, PA, USA; Department of Neurobiology, University of Pittsburgh, Pittsburgh, PA, USA; Department of Neurosurgery, Graduate School of Medical Sciences, Kyushu University, Fukuoka, Japan; Children’s Medical Center Research Institute, University of Texas Southwestern Medical Center, Dallas, TX, USA; Harold C. Simmons Comprehensive Cancer Center, University of Texas Southwestern Medical Center; Dallas, TX, USA; Department of Pediatrics, University of Texas Southwestern Medical Center, Dallas, TX, USA; Peter O’Donnell Jr. Brain Institute, University of Texas Southwestern Medical Center, Dallas, TX, USA

## Abstract

Glioblastoma (GBM) stem-like cells (GSCs) contribute to therapeutic resistance and recurrence. We sought to define cellular processes underlying GSC resilience. We discovered that GSCs, unlike differentiated GBM cells (DGCs) or non-malignant neural cells, depend on mitochondrial function for survival. To target this vulnerability, we exploited doxycycline (DOXY), an antibiotic used in humans, to interfere with mitochondrial protein translation. DOXY induced cell death and inhibited sphere formation in GSCs, but not in DGCs or non-malignant cells, indicating a differentiation state-selective effect. Mechanistically, DOXY induced mitochondrial dysfunction and activated a stress-responsive apoptotic program involving HRI-mediated signaling. DOXY displayed antitumor efficacy in patient-derived GBM organoid and orthotopic xenograft models. Our study reveals that DOXY can selectively target undifferentiated glioma cells, informing a drug repurposing-based strategy.

## INTRODUCTION

Glioblastoma (GBM) is the most aggressive form of malignant brain tumor and remains one of the most lethal human cancers, with an extremely poor prognosis*(1)*. The current standard of care consists of maximal surgical resection followed by radiotherapy and temozolomide (TMZ) chemotherapy*(2)*. However, GBM almost inevitably develops therapeutic resistance, thought to be driven principally by GBM stem-like cells (GSCs)*(3)*. The cancer stem cell concept, initially established in hematologic malignancies, was also found to be highly relevant in GBM, where a stem-like subpopulation was shown to possess tumor-initiating capacity and self-renewal*(4)*. Large-scale genomic analyses have further revealed that GBM exhibits profound cellular heterogeneity*(5–8)*, which limits the efficacy of uniform therapeutic approaches. Importantly, GSCs preferentially survive radiotherapy through enhanced DNA damage response and repair mechanisms and are maintained in vivo by specialized tumor microenvironments such as the perivascular niche*(9, 10)*. Consequently, effective targeting of GSCs remains a central yet unresolved translational challenge in the treatment of GBM. Treatment strategies that preferentially eradicate GSCs may mitigate treatment resistance mechanisms and improve patient outcomes.

Mitochondria play central roles in cellular metabolism and mitochondrial dysfunction is implicated in various diseases, including cancer*(11)*. In human GBM, in vivo ^13^C tracing and magnetic resonance spectroscopy studies have demonstrated that tumors retain substantial mitochondrial oxidative metabolism of glucose-derived carbon in patients*(12, 13)*. Consistent with these human data, GBM patient-derived xenograft models display high mitochondrial respiration and a marked dependence on oxidative phosphorylation, and mitochondrial metabolic pathways can represent actionable therapeutic vulnerabilities in vivo*(14, 15)*.

Mitochondrial translation has been explored as a potential target via the use of antibiotics as anticancer agents*(16)*. Because many antibiotics are non-toxic and clinically approved, repurposing them for cancer therapy could circumvent barriers to clinical translation that can impede the development of experimental therapeutics. Matsumoto et al. demonstrated that doxycycline (DOXY), a tetracycline antibiotic, inhibits mitochondrial translation, induces mitochondrial dysfunction, and triggers endoplasmic reticulum (ER) stress-dependent apoptosis through mitochondria-ER crosstalk*(17)* in undifferentiated prostate cancer cells. Other studies have shown that electron transport chain inhibitors such as oligomycin cause mitochondrial dysfunction and upregulate expression of the integrated stress response (ISR) pathway effector ATF4 through activation of the OMA1-DELE1-HRI-p-eIF2α signaling axis in non-tumor cells*(18)*. These findings position mitochondrial stress as a direct upstream regulator of the ISR and ATF4-dependent cellular gene expression programs. Although DOXY has been reported to be effective against certain cancer or stem cells*(19, 20)*, the precise mechanism underlying DOXY-induced cell death remains unclear, and its efficacy against GSCs and its therapeutic potential in glioma have not yet been fully elucidated.

Here, we used gene expression profiling and oxygen consumption assays to investigate whether GSCs and DGCs display differential reliance on mitochondrial function. We found that GSCs display elevated mitochondrial metabolism and, further, that inhibiting mitochondrial translation via treatment with DOXY, an antibiotic in wide clinical use, selectively eliminated GSCs. Our study reveals a previously unrecognized mechanism by which DOXY disrupts mitochondrial homeostasis and triggers stress-induced apoptosis in GSCs, highlighting a potential therapeutic strategy to selectively target GSC populations in GBM.

## RESULTS

### GSCs exhibit elevated mitochondrial activity compared to DGCs

To examine differences in mitochondrial dependence between glioma cell states, we leveraged transcriptomic and mitochondrial activity data collected from isogenic pairs of GSCs and DGCs (Fig. 1A). DGCs were generated from GSCs by culturing them under serum-replete DMEM medium. We first assessed the mitochondrial metabolism-related gene expression signatures in RNA-seq datasets*(21)*. GSEA confirmed significant enrichment of mitochondrial pathways including oxidative phosphorylation (OXPHOS), trycarboxylic acid (TCA) cycle, and reverse electron transport (RET) pathways in GSCs (Fig. 1B-D). Supporting this observation, gene ontology (GO) enrichment analysis further identified an upregulation of mitochondria-associated biological processes and cellular components in GSCs (fig. S1A).

**Fig. 1.**
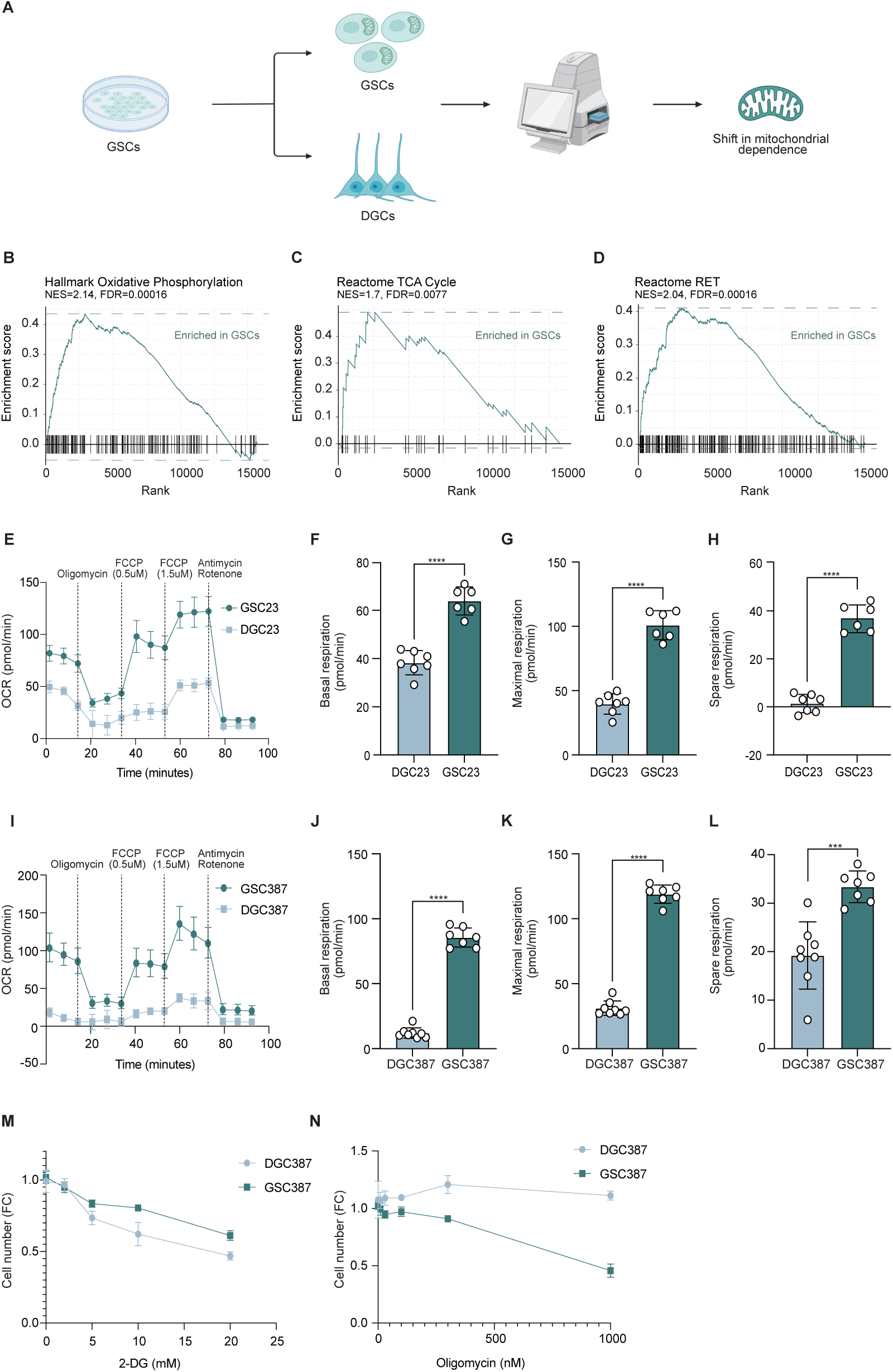
GSCs are dependent on mitochondrial metabolism. (A) Schematic overview of the experimental design. DGCs were generated from GSCs by culturing under serum-containing (DMEM) conditions, and the metabolic shift in mitochondrial dependence was compared. (B-D) Gene set enrichment analysis (GSEA) showing enrichment of oxidative phosphorylation (from Hallmark, B), the TCA cycle (C), and RET pathways (D) (from Reactome) in GSCs. (E) Oxygen consumption rate (OCR) profiles from mitochondrial stress tests in GSC23 and DGC23 (GSC23, n = 7; DGC23, n = 6). (F-H) Quantification of basal respiration (F), maximal respiration (G), and spare respiratory capacity (H). (I) OCR profiles from mitochondrial stress tests in GSC387 and DGC387 (GSC387, n = 8; DGC387, n = 7). (J-L) Quantification of basal respiration (J), maximal respiration (K), and spare respiratory capacity (L). (M-N) Cell viability (n=4) following treatment with glycolysis inhibitor 2-deoxy-D-glucose (2-DG) (M) or the mitochondrial inhibitor oligomycin (N). Data are presented as mean ± SD. Statistical analyses were performed using an unpaired two-tailed Student’s t-test (F-H, J-L). \*\*\**p* < 0.001; \*\*\*\**p* < 0.0001.

To validate transcriptomic differences at the protein level, we compared mitochondrial protein expression between GSCs, DGCs, and non-malignant neural cells (including NHA immortalized astrocytes and NSC11 neural stem cells). Among mitochondrial proteins, we focused on MT-CO1 and MT-CO2, components of Complex IV in the mitochondrial respiratory chain. These proteins are encoded by genes in the mitochondrial genome and translated via mitochondrial ribosomes. Both MT-CO1 and MT-CO2 were higher in GSCs compared with non-malignant cells and DGCs across multiple lines (fig. S1B). In an isogenic comparison, MT-CO2 was elevated in GSC387 relative to DGC387 (fig. S1C-D). Therefore, at both transcriptomic and protein levels, GSCs consistently showed elevated expression of mitochondria-related genes and proteins.

Next, we performed Seahorse assays to measure mitochondrial activity using oxygen consumption rate (OCR) as a proxy. All GSC lines displayed substantially higher OCR values than DGCs, indicating increased mitochondrial respiration (Fig. 1E-L, fig. S1E). Consistently, tetramethylrhodamine ethyl ester (TMRE) staining, which reflects mitochondrial membrane potential and serves as an indicator of mitochondrial activity*(22)*, revealed higher mitochondrial membrane potential in GSCs compared with DGCs (fig. S1F-H). To determine whether GSCs are more dependent on mitochondrial metabolism than DGCs, we examined the effects of blocking either glycolysis or mitochondrial OXPHOS. Treatment with 2-deoxyglucose (2-DG), a glycolysis inhibitor*(23)*, and oligomycin, a mitochondrial ATP synthase inhibitor*(24)*, allowed us to assess the relative dependency of each cell type. GSCs were less sensitive to 2-DG but more sensitive to oligomycin than DGCs (Fig. 1M and N), indicating a stronger reliance on mitochondrial oxidative metabolism. This phenotype may reflect the unique bioenergetic demands of GSCs*(25, 26)*, which rely on oxidative metabolism to sustain stemness and resist metabolic stress. To assess whether undifferentiated cell states are linked to elevated mitochondrial metabolism across tumor types, we analyzed public RNA-seq dataset of ovarian cancers (GSE148003)*(27)*. Genes upregulated in cancer stem cells (CSCs) relative to differentiated cells showed significant enrichment of the GO term mitochondrial gene maintenance (fig. S1I). These results suggest that enhanced mitochondrial activity and metabolism in undifferentiated tumor cells are observed across cancer types.

### Inhibiting mitochondrial translation with doxycycline selectively targets GSCs

We sought to identify drugs that induce mitochondrial dysfunction and could be used to target the heightened dependence of GSCs on mitochondrial function. Because mitochondria evolved from bacteria, many antibiotics that target bacterial ribosomes can also impair mitochondrial translation. Importantly, these antibiotics typically affect only mitochondria or bacteria, without harming translation of transcripts from nuclear genes in eukaryotic cells*(28)* (Fig. 2A). Among such agents, we focused on clinically approved drugs that can penetrate the blood-brain barrier (BBB).

**Fig. 2.**
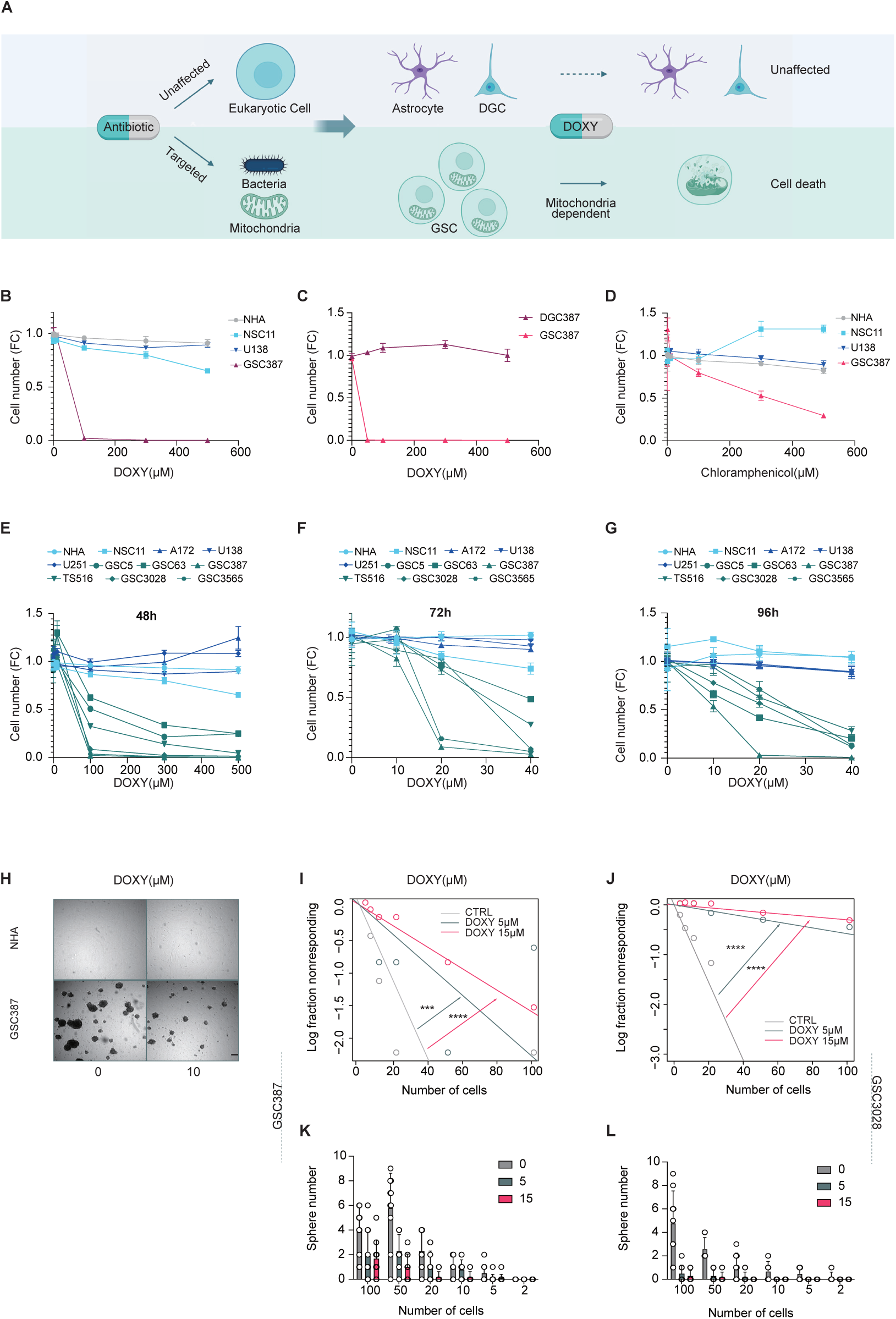
Doxycycline selectively targets GSCs, inhibiting sphere formation and inducing cell death. (A) Schematic illustration of the hypothesis. Antibiotics specifically target mitochondria and are therefore safe for normal human cells. Because GSCs are highly dependent on mitochondrial function, mitochondrial inhibition is expected to selectively affect GSCs. (B, E) represent the same experimental set. (B) Representative cell viability after 48 h of treatment with doxycycline (DOXY) (n = 3). (C) Cell viability after 24 h of DOXY treatment between GSC387 and DGC387 (n = 3). (D) Representative cell viability after 72 h of treatment with chloramphenicol (n = 3). (E) Cell viability after 48h of treatment with DOXY (n = 3-4). (F, G) Cell viability after 72, and 96 h of treatment with DOXY (n = 3-4). (H) Representative images of NHA and GSC387 treated with 10 μM DOXY for 96 h. Scale bar, 100 μm. (I-L) Extreme limiting dilution assay (H, I) and sphere-formation assay (> 50 μm) (J, K) of GSC387 and GSC3028 treated with control, 5 μM, or 15 μM DOXY (n=10). Data are presented as mean ± SD. Statistical analysis was performed using one-way ANOVA by Dunnett’s multiple comparisons test. \*\*\**p* < 0.001; \*\*\*\**p* < 0.0001.

We evaluated the effects of several mitochondrial translation inhibitors, including DOXY, chloramphenicol, and linezolid, on neural cell viability*(29–31)*. Initial 48-hour DOXY treatment revealed selective GSC death at lower doses (Fig. 2B). To determine whether this selective vulnerability was evident even at earlier time points, we next examined the effects of 24-hour DOXY treatment in an isogenic pair of glioma cells (GSC387 and DGC387). DOXY selectively affected GSC387, while having little impact on DGC387 (Fig. 2C). By contrast, 72-hour treatment with chloramphenicol or 48-hour treatment with linezolid did not produce comparable cytotoxicity. Chloramphenicol showed some cytotoxicity toward GSCs but only at relatively high concentrations and linezolid did not induce apparent cell death even at high doses. (Fig. 2D, fig. S2A and B). Importantly, acutely supplementing GSCs with FBS or pyruvate (two components that differ substantially between GSC and DGC media) did not rescue sensitivity of GSCs to DOXY (fig. S2C and D).

Next, we first examined the effects of 48 h DOXY treatment on cell viability to establish a broad dose-response relationship and confirm selective sensitivity of GSCs (Fig. 2E). We then assessed the effects of prolonged, lower-dose exposure (72-96 h) (Fig. 2F and 2G), which revealed substantial GSC death; for example, GSC387 exhibited approximately 50% loss of viability after 96 h of treatment with only 10 µM DOXY. This concentration also reduced sphere size and integrity (Fig. 2H). To further assess whether DOXY inhibits self-renewal capacity, we performed an extreme limiting dilution assay (ELDA). Even at 5 µM, DOXY markedly suppressed sphere formation of GSC387 and GSC3028 lines, indicating a strong inhibition of self-renewal (Fig. 2I-L). Although sensitivity to DOXY varied moderately among GSC lines, 20 μM DOXY potently inhibited sphere formation in all lines (fig. S2E-H). Considering that DOXY accumulates to 0.9-5.6 μM in CSF after standard dosing (200 mg/day in humans) *(32)*, these values provide a clinically relevant reference range for interpreting the low-dose effects observed in prolonged in vitro exposure experiments. Finally, cell death assays revealed that DOXY treatment increased the proportion of apoptotic GSC387 and GSC3028 cells (fig. S2I-J). Considering these findings together with the ability of DOXY to cross the BBB, its effectiveness at low micromolar concentrations, and its well-established clinical safety profile, DOXY may display antitumor activity against glioma by targeting GSCs in preclinical models.

### Doxycycline induces mitochondrial dysfunction, ROS accumulation, and apoptosis in GSCs

We next investigated the mechanism by which DOXY induces cell death in GSCs. To delineate the mechanisms underlying DOXY-induced GSC death, subsequent mechanistic experiments employed higher DOXY concentrations over shorter time courses to robustly induce mitochondrial stress and enable pathway interrogation. In contrast, prolonged exposure to low micromolar DOXY concentrations, within the range observed in human cerebrospinal fluid, was sufficient to suppress GSC self-renewal and sphere formation. We hypothesized that DOXY causes mitochondrial dysfunction, leading to a loss of mitochondrial membrane potential and accumulation of fragmented mitochondria (Fig. 3A). Following DOXY treatment, expression of the mitochondrially encoded MT-CO1 protein, a component of Complex IV, was reduced, consistent with impaired mitochondrial translation (Fig. 3B). In parallel, DOXY increased the expression of the mitochondrial damage marker GDF15 (Fig. 3C). These results demonstrate that DOXY induces mitochondrial dysfunction and mitochondrial damage.

**Fig. 3.**
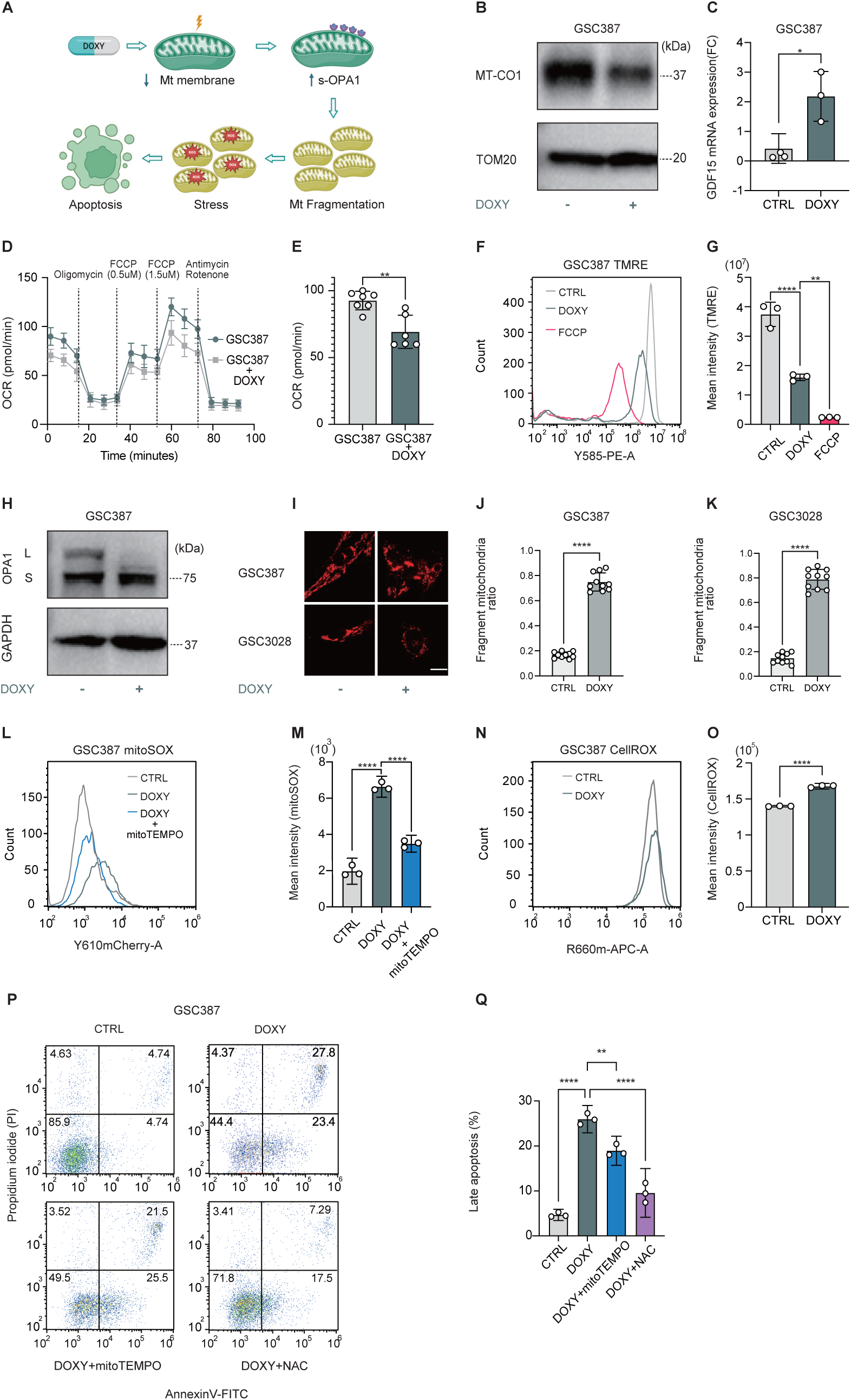
Doxycycline inhibits mitochondrial translation, leading to decreased mitochondrial membrane potential and generation of reactive oxygen species (ROS). DOXY concentrations were selected for mechanistic dissection; unless indicated, DOXY was used at 30 μM. (A) Schematic illustration of loss of mitochondrial membrane potential, increased s-OPA1, and mitochondrial fragmentation. (B-C) MT-CO1 protein and GDF15 mRNA expression after 24h treatment (n = 3). (D-E) OCR curves and maximal respiration in GSC387 cells after 2h treatment (n = 6-7). (F-G) TMRE flow cytometry in control, DOXY, or FCCP (10 μM) treated cells (3 h) (n = 3). (H) OPA1 protein expression after 24h treatment (n=3). (I) TOM20 immunofluorescence in GSC387 and GSC3028 (60 μM) cells after 24 h treatment. Scale bar, 10 μm. (J-K) Quantification of mitochondrial fragmentation (n = 10). (L-M) MitoSOX flow cytometery in control, DOXY, and DOXY + mitoTEMPO (20 μM)-treated cells after 3 h (n=3). (N-O) CellROX flow cytometry in control and DOXY (30 μM) treated cells after 3 h (n=3). (P-Q) Apoptosis analysis and quantification in control, DOXY (30 μM), DOXY+ mitoTEMPO (30 μM), and DOXY + NAC (0.5 mM) treated cells after 24h (n=3). Data are mean ± SD. Statistics used unpaired Student’s *t*-test or one-way ANOVA with Sidak’s test. \**p* < 0.05; \*\**p* < 0.01; \*\*\**p* < 0.001; \*\*\*\**p* < 0.0001.

To further assess the functional consequences of this dysfunction, we examined whether DOXY alters mitochondrial activity, membrane potential, and morphology, as mitochondrial shape reflects functional status*(33)*. As early as 2-3 hours after treatment, DOXY decreased OXPHOS activity as measured by Seahorse assay and reduced mitochondrial membrane potential as assessed by TMRE staining (Fig. 3D-G, fig. S3A). We next examined proteins associated with mitochondrial morphology and integrity. Consistent with mitochondrial stress, DOXY increased the ratio of the short (s-OPA1) to long (l-OPA1) forms of OPA1 (Fig. 3H), indicating proteolytic cleavage of the OPA1 protein. OPA1 proteolysis promotes mitochondrial fragmentation*(34, 35)*. Consistent with this increase in OPA1 processing, the proportion of fragmented mitochondria was significantly increased by DOXY treatment (Fig. 3I-K, fig. S3B-C). Others have reported that oxidative stress is associated with mitochondrial fragmentation*(36)*. Therefore, we hypothesized that DOXY induces mitochondrial dysfunction and fragmentation which in turn lead to elevated ROS production, likely resulting from impaired electron transport and enhanced superoxide generation. To test this, we measured mitochondrial and cytosolic ROS levels using MitoSOX and CellROX, respectively. DOXY markedly increased MitoSOX fluorescence, which was attenuated by the mitochondrial ROS scavenger mitoTEMPO (Fig. 3L-M). CellROX staining also revealed a modest increase in cytosolic ROS following DOXY treatment (Fig. 3N-O). Apoptosis assays confirmed that DOXY increased the proportion of apoptotic cells, whereas co-treatment with N-acetylcysteine (NAC), a general ROS scavenger, or mitoTEMPO significantly reduced apoptosis, supporting a role for oxidative stress in this response (Fig. 3P-Q). To determine whether ROS directly triggers apoptosis, cells were treated with rotenone (a mitochondrial complex I inhibitor that generates superoxide) or hydrogen peroxide (H₂O₂), which induces cytosolic ROS. To confirm that ROS mediated stress activates downstream apoptosis signaling, we also examined the expression of ATF4, stress responsive transcription factor*(37)*. Both rotenone and H₂O₂ have been reported to increase ROS and cause apoptosis via ATF4 activation*(38, 39)*. Consistent with these reports, both agents increased ATF4 expression and enhanced apoptosis (fig. S3D-H). Conversely, NAC attenuated H₂O₂-induced apoptosis, and mitoTEMPO rescued the cell death caused by rotenone (fig. S3G-H). Together, these findings demonstrate that DOXY-induced mitochondrial dysfunction leads to mitochondrial fragmentation and promotes ROS accumulation, which in turn activates apoptotic signaling in GSCs.

### DOXY induces apoptosis through activation of the HRI-p-eIF2α-ATF4-CHOP-TRIB3 axis

We next investigated the mechanism by which DOXY induces apoptosis following ROS production and mitochondrial fragmentation. We hypothesized that, in addition to ROS generation, another major pathway contributes to DOXY-induced apoptosis, as the rescue effects of mitoTEMPO and NAC were only partial. We speculated that, upon mitochondrial fragmentation, two parallel activations occur, one through the OMA1-OPA1-DELE1 pathway and the other through ROS accumulation, both converging on the activation of the HRI-p-eIF2α-ATF4-CHOP-TRIB3 axis, which mediates this apoptotic response (Fig. 4A).

**Fig. 4.**
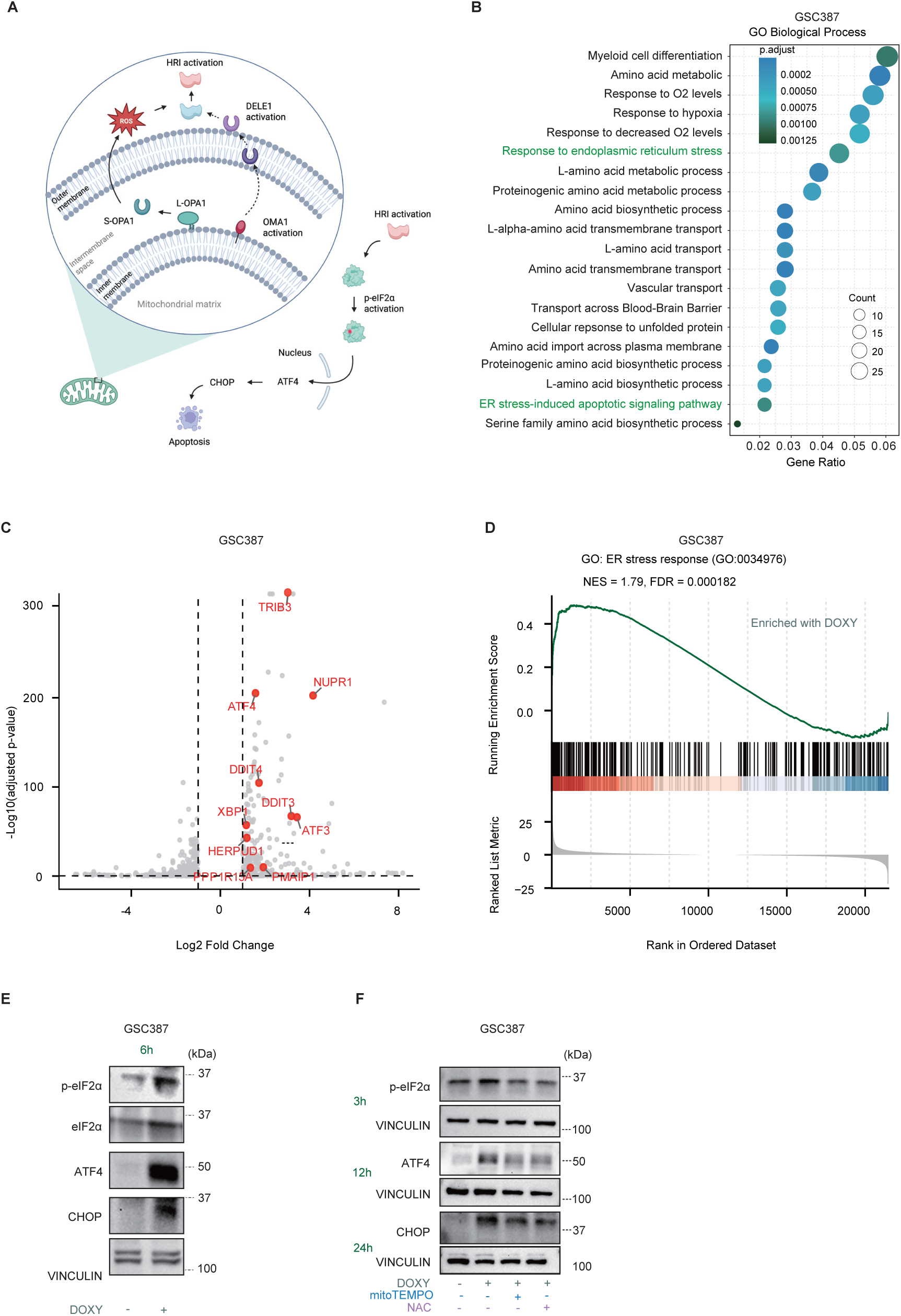
Doxycycline activates the integrated stress response (ISR) in GSCs. (A) Schematic illustration showing that DOXY activates OMA1, leading to increased s-OPA1 and DELE1 cleavage. Activated DELE1 subsequently stimulates HRI, which triggers phosphorylation of eIF2α and activation of the ATF4-CHOP-TRIB3 apoptotic pathway. (B) Gene Ontology (GO) analysis of RNA-seq data from GSC387 treated with or without DOXY (20 μM, 24 h). (C) Volcano plot of differentially expressed genes in GSC387 with or without DOXY treatment. (D) GSEA showing enrichment of endoplasmic reticulum (ER) stress-related pathways (GO:0034976) after DOXY treatment. (E, F) Representative western blots showing expression of ISR under the indicated conditions. (E) p-eIF2α, eIF2α, ATF4, and CHOP expression in GSC387 with or without DOXY (30 μM, 6 h) (n=3). (F) Time-course effects of ROS scavengers (mitoTEMPO (20 μM), NAC (0.5 mM)) on DOXY-induced ISR activation (n=3).

To explore this mechanism, we performed RNA-seq analysis of GSC387 or TS516 cells treated with or without DOXY. GO analysis revealed significant enrichment of ER stress-related pathways, while Reactome pathway analysis indicated upregulation of HRI-mediated and ATF4-dependent signaling in both GSC lines (Fig. 4B, fig. S4A-C). Volcano plots also showed elevated expression of genes belonging to the ATF4-CHOP-TRIB3 axis (Fig. 4C, fig. S4D), and GSEA confirmed the enrichment of ER stress-response signatures following DOXY administration (Fig. 4D).

At the protein level, DOXY treatment increased the levels of p-eIF2α, ATF4, and CHOP, with CHOP induction detectable after 6 hours of treatment, but not after 3 hours (Fig. 4E, fig. S4E). Time-course experiments further showed that both mitoTEMPO and NAC reduced p-eIF2α, ATF4, and CHOP expression, implying that ROS is involved in the activation of this pathway (Fig. 4F).

Because OPA1 processing is regulated by OMA1 activation*(18, 40)*, we hypothesized that the increased ratio of s-OPA1 to l-OPA1 was driven by OMA1. Indeed, OMA1 knockdown (shOMA1) reduced this ratio during DOXY treatment (Fig. 5A, fig. S5A and B). Previous reports have demonstrated that OMA1-mediated DELE1 cleavage activates HRI*(18)*. Our Reactome analysis suggested DOXY activates HRI, raising the possibility that OMA1 and DELE1 are involved in this response. Consistent with this, DOXY treatment was associated with increased DELE1 cleavage (Fig. 5B). To further evaluate this pathway, we used hemin to block HRI activation. As expected, DOXY increased p-eIF2α and ATF4 expression, while hemin treatment attenuated both (Fig. 5C).

**Fig. 5.**
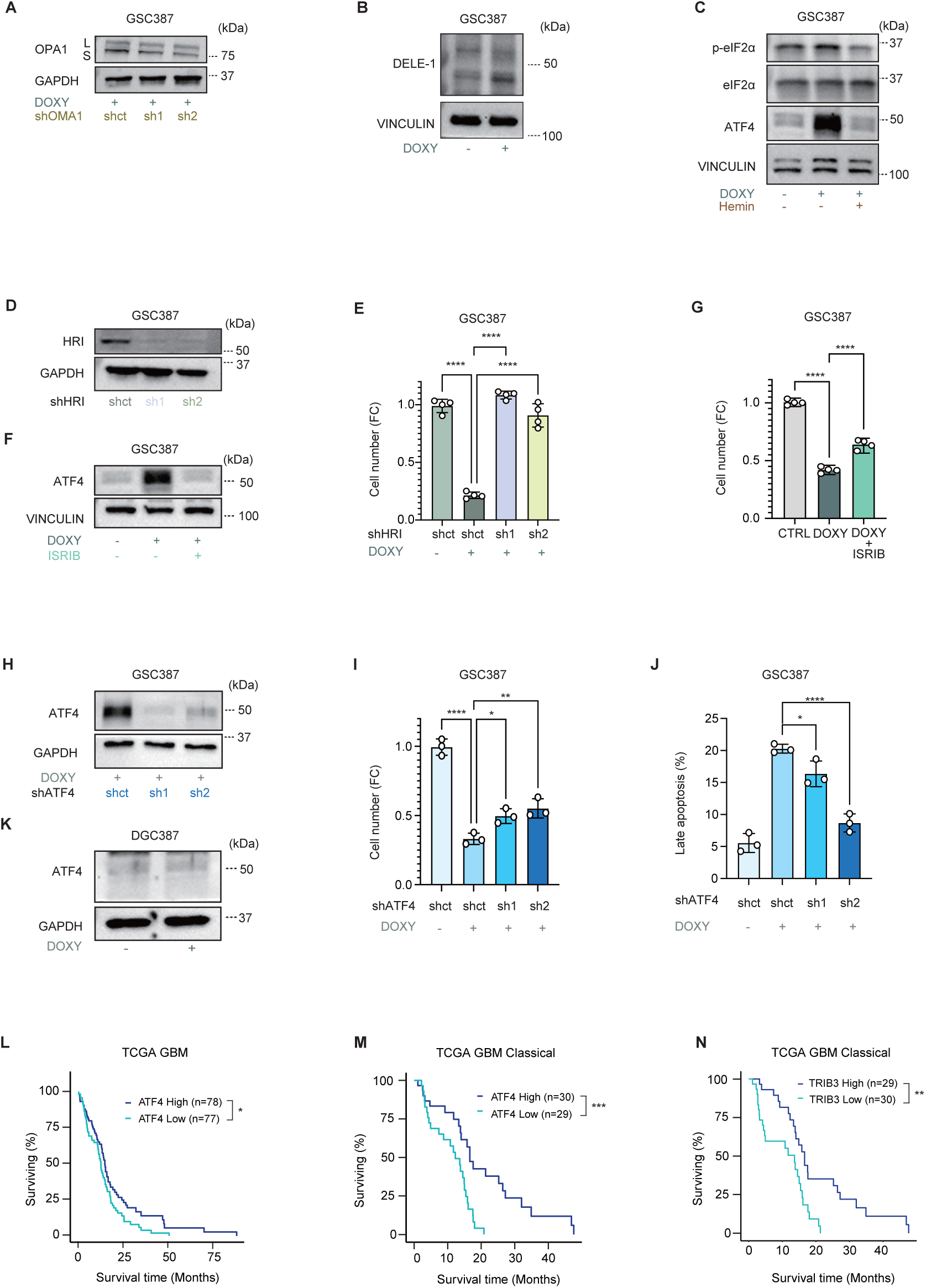
Doxycycline-induced ISR activation triggers apoptosis through the HRI-p-eIF2α-ATF4-CHOP-TRIB3 axis. (A) Effects of shOMA1 on DOXY-induced signaling (30 μM, 3 h; n = 3). (B) DELE1 cleavage with DOXY (30 μM, 24 h; n= 3). (C) p-eIF2α and ATF4 expression in GSC387 treated with DOXY (30 μM) and/or hemin (30 μM; n= 3). (D) Knockdown efficiency of shHRI. (E) Cell viability of GSC387 treated with DOXY (40 μM, 24 h) with or without shHRI (n = 4). (F) p-eIF2α and ATF4 expression in GSC387 treated with DOXY (30 μM, 24 h) and ISRIB (2 μM; n= 3). (G) Cell viability of GSC387 treated with DOXY (30 μM, 24 h) and ISRIB (1 μM; n= 4). (H-J) Effects of shATF4 on ATF4 expression (H; 3 h), Cell viability (I; 24 h), and apoptosis (J; 24 h) in DOXY-treated GSC387 (30 μM; n= 3). (K) ATF4 expression in DGC387 with or without DOXY (30 μM, 3 h). (L-N) Kaplan-Meier survival analyses of TCGA GBM cohorts stratified by ATF4 expression(L) and, within the Classical subtype, by ATF4 (M) or TRIB3 (N) expression. Data are presented as mean ± SD. Statistical analyses were performed using one-way ANOVA with Sidak’s multiple comparisons test. \**p* < 0.05; \*\**p* < 0.01; \*\*\**p* < 0.001; \*\*\*\**p* < 0.0001.

We further interrogated HRI as a downstream effector of DOXY treatment via genetic knockdown experiments. Cells in which HRI was knocked down (shHRI) exhibited reduced HRI expression and were almost completely rescued from DOXY-induced cytotoxicity (Fig. 5D-E). We then examined other key mediators of the ISR to determine the influence of ISR and ATF4 pathways. Both ATF4 knockdown (shATF4) and treatment with ISRIB, a selective ISR inhibitor, decreased ATF4 expression and partially rescued DOXY-induced cell death, although their effects were weaker than those of HRI knockdown (Fig. 5F-I, fig. S5C-F). Apoptosis assays further confirmed that shATF4 reduced DOXY-induced apoptosis (Fig. 5J, fig. S5D).

These findings position HRI as a central mediator linking mitochondrial stress to the ISR. These data suggest that DOXY treatment specifically activates the recently described OMA1-DELE1-HRI signaling axis in GSCs. This pathway enables cells to sense and respond to mitochondrial dysfunction*(18)*. We next examined the downstream effectors of ATF4, including CHOP and TRIB3. Both CHOP and TRIB3 knockdown significantly restored GSC fitness during DOXY treatment (fig. S5G-J).

To assess the specificity of HRI activation for the ISR response triggered by DOXY, we inhibited the four kinases known to act upstream of p-eIF2α (HRI, PKR, PERK, and GCN2)*(41)*. PKR and PERK inhibitors (PRK-IN-C16 and GSK2656157) unexpectedly increased ATF4 expression, whereas GCN2 inhibition (GCN2iB) modestly reduced it (fig. S5K-M). However, the extent of reduction was smaller than that achieved by hemin, supporting the conclusion that HRI is the primary kinase involved in DOXY’s mechanism of action. Collectively, these findings establish a direct mechanistic link between mitochondrial dysfunction and stress-responsive apoptosis, positioning the HRI-eIF2α-ATF4 pathway as a pivotal mediator of mitochondrial stress signaling in GSCs treated with DOXY.

The effect of DOXY on this pathway was specific to GSCs because we did not observe increased ATF4 expression in DGCs, including DGC387 and A172 cells, under the same conditions (Fig. 5K, fig. S5N). These findings complement RNA-seq results showing that GSCs exhibit higher baseline ATF4 expression compared with DGCs (fig. S5O) and suggest that GSCs may be uniquely susceptible to stress-induced ATF4 upregulation. OPA1 expression was also significantly higher in GSCs than in DGCs (fig. S5P). Clinically, lower ATF4 expression is associated with poor prognosis in GBM patients. When restricted to the Classical subtype, lower ATF4 and TRIB3 expression are both associated with poor prognosis (Fig. 5L-N). Therefore, ISR activation may be generally tumor suppressive, although more work is needed to understand this correlative relationship.

### Disruption of an adaptive mitophagy response potentiates DOXY-induced GSC killing

We next explored how to enhance the antitumor effect of DOXY for potential clinical application. Given our findings that DOXY induces mitochondrial damage, we hypothesized that DOXY treatment would increase the number of damaged mitochondria, leading to elevated ROS levels. To counteract this stress, cells may engage an adaptive mitochondrial quality control program, including mitophagy as a cytoprotective mechanism. Therefore, we reasoned that mitophagy represents an adaptive response to DOXY-induced mitochondrial stress, and that disruption of this adaptive response would exacerbate mitochondrial dysfunction and ROS accumulation, thereby promoting cell death (Fig. 6A).

**Fig. 6.**
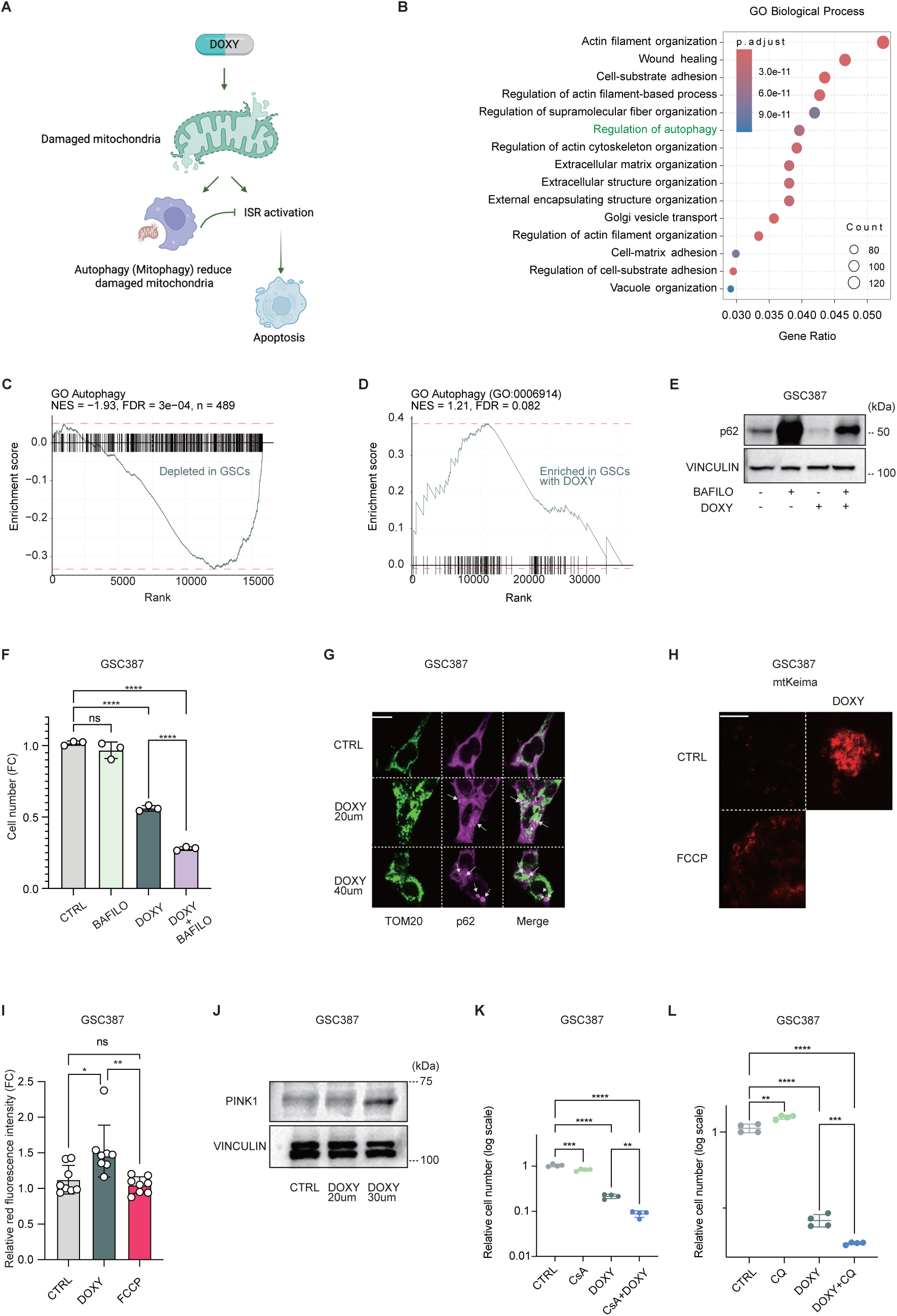
Mitophagy protects GSCs against doxycycline, and inhibition of mitophagy enhances the cytotoxic effect of DOXY. (A) Schematic illustrating DOXY-induced mitochondrial dysfunction and mitophagy activation. (B-D) GO/GSEA analyses showing enrichment of autophagy pathways in GSCs vs DGCs and in GSC387 after DOXY. (E) Western blot analysis of GSC387 treated with DOXY (20 μM) and bafilomycin (BAFILO) (20 nM) for 24 h (n=3). (F) Cell viability of GSC387 treated with or without DOXY (30 μM) and BAFILO (20 nM) for 24 h (n=3). (G) Immunofluorescence images of TOM20 and p62 in GSC387 treated with DOXY (20 μM, 40 μM) for 24 h. Scale bar: 10 μm. (H-I) mt-Keima imaging in GSC387 under control, DOXY (50 μM, 12 h), or FCCP (10 μM, 3 h) conditions and quantification of red fluorescence (n=9) Scale bar: 50 μm. (J) PINK1 protein expression in GSC387 after DOXY (20 μM, 30 μM) for 24 h. (K-L) Cell viability in GSC387 treated with DOXY (30 μM) ±CsA (2 μM) or chloroquine (CQ, 10 μM) for 24 h (n=4). Data are mean ± SD. Statistical analyses were performed using one-way ANOVA followed by Sidak’s multiple comparisons test. \**p* < 0.05; \*\**p* < 0.01; \*\*\**p* < 0.001; \*\*\*\**p* < 0.0001.

Because DOXY was less effective in DGCs than in GSCs, we compared transcriptomic profiles of these two cell types to identify possible differences that could explain this phenomenon. Analysis of RNA-seq datasets comparing GSCs and DGCs revealed that autophagy-related pathways were differentially regulated (Fig. 6B). GSEA further demonstrated significant enrichment of autophagy-related gene sets in DGCs compared with GSCs (Fig. 6C). Moreover, DOXY treatment further enhanced autophagy-related gene sets enrichment (Fig. 6D).

These findings support our hypothesis that autophagy or mitophagy plays a protective role under DOXY-induced stress. Because GSCs have lower expression of autophagy- and mitophagy-related genes, they may be more sensitive to mitochondrial damage. To test this hypothesis, we next examined whether DOXY affects autophagic flux. DOXY treatment altered the abundance of the autophagy adaptor protein p62, whereas the autophagy inhibitor bafilomycin increased it (Fig. 6E). Co-treatment with DOXY and bafilomycin induced greater killing of GSC387 and TS516 GSC lines relative to DOXY treatment alone (Fig. 6F, fig. S6A).

To visualize autophagic structures, we assessed the localization of TOM20 and p62 by immunofluorescence. After DOXY administration, p62 formed numerous cytoplasmic puncta, some of which colocalized with mitochondria, suggesting recruitment of damaged mitochondria to autophagy-related structures (Fig. 6G). Together, these results indicate that DOXY induces autophagy and mitophagy as a cytoprotective adaptive response, and that inhibiting this response enhances the cytotoxic effect of DOXY.

To directly quantify mitophagy, we utilized mt-Keima, a pH-sensitive fluorescent reporter that shifts from green to red upon lysosomal delivery of mitochondria*(42)*. DOXY treatment markedly increased red fluorescence intensity, reflecting enhanced mitophagy (Fig. 6H-I). Flow cytometric analysis further confirmed an increase in the proportion of mitophagy-positive cells following DOXY exposure (fig. S6B). Consistent with these findings, protein analysis revealed upregulation of PINK1 after treatment with moderate doses of DOXY (Fig. 6J), suggesting activation of the PINK1-Parkin mitophagy pathway.

To determine the functional relevance of DOXY-induced mitophagy, having already confirmed the effect of bafilomycin, we next evaluated additional pharmacological modulators of mitochondrial stress response, including cyclosporin A (CsA)*(43)* and P110, a selective inhibitor of Drp-1-Fis1 interaction that suppresses mitochondrial fission and can indirectly influence mitophagy*(44)*. Co-treatment with DOXY and CsA increased cell death compared with DOXY alone (Fig. 6K). P110 also enhanced DOXY-induced killing under conditions in which DOXY alone did not induce maximal cytotoxicity (fig. S6C).

Finally, considering the clinical limitations of bafilomycin, we evaluated chloroquine (CQ) as a safer autophagy inhibitor. Similar to bafilomycin, CQ enhanced DOXY-induced GSC killing (Fig. 6L). Together, these findings demonstrate that DOXY induces a cytoprotective mitochondrial adaptive response associated with engagement of mitophagy, and that disruption of this adaptive program-particularly through inhibition of autophagic flux-can potentiate the cytotoxic effect of DOXY in GSCs. Accordingly, combination therapy involving DOXY and agents that interfere with mitochondrial stress adaptation, including autophagy or mitophagy, may represent a promising strategy to potentiate the antitumor activity of DOXY.

### Doxycycline displays therapeutic efficacy in patient-derived organoid and xenograft models

We next evaluated the therapeutic potential of DOXY in organoid and orthotopic xenograft models of GBM. Surgically explanted organoids (SXOs) were created from a human GBM specimen*(45, 46)* and treated with DOXY, followed by histological and immunohistochemical analyses. Hematoxylin-eosin (H&E) staining revealed that DOXY reduced cell density in the central region of the organoids, while immunostaining showed decreased SOX2 expression and increased cleaved caspase-3 levels (Fig. 7A-C). These findings indicate that DOXY effectively suppresses stemness and induces apoptosis in the tumor microenvironment.

**Fig. 7.**
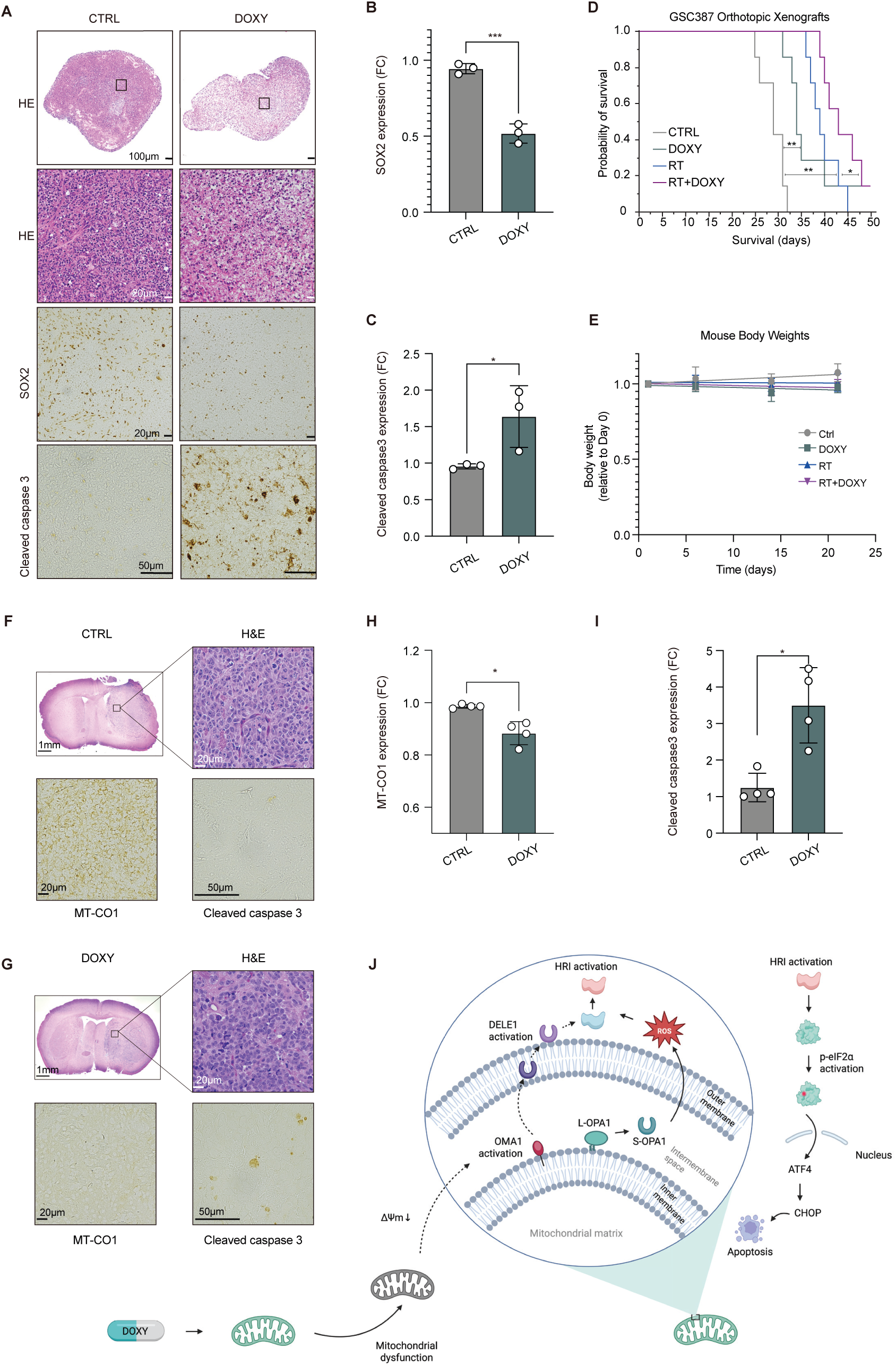
Doxycycline demonstrates therapeutic efficacy in organoid and xenograft GBM models. (A) Representative images of patient-derived SXO organoids treated with DOXY (35 μM, 96 h) stained with H&E, SOX2, and cleaved caspase-3. Scale bar, 100 μm; H&E/SOX2 (20×), 20 μm; cleaved caspase-3 (50×), 50 μm. (B) Quantification of SOX2-positive cells. (C) Quantification of cleaved caspase-3-positive cells. (D) Kaplan-Meier survival curves of mice bearing GSC387 orthotopic xenografts (n = 7). (E) Body weight changes in mice treated with control, DOXY (40 mg/kg/day), RT (2 Gy × 3), or RT + DOXY. (F-G) Representative H&E, MT-CO1, and cleaved caspase 3 immunostaining in (F) control and (G) DOXY-treated tumors. Scale bar, 1 mm or 10 μm. (H) Quantification of SOX2-positive cells. (I) Quantification of cleaved caspase-3-positive cells. (J) Schematic summarizing the proposed mechanism of DOXY action. Data are presented as mean ± SD. Statistical analyses between two groups were performed using an unpaired two-tailed Student’s *t*-test (B, C) or using the Mann-Whitney U test (H, I). Survival curved for four groups were analyzed using the Kaplan-Meier method and compared by the log-rank test (D). \**p* < 0.05; \*\**p* < 0.01; \*\*\**p* < 0.001.

We then examined the efficacy of DOXY in an orthotopic xenograft model. To maintain clinical relevance, we used a non-toxic, human-equivalent dose of DOXY in mice. Because radiotherapy (RT) is a standard treatment for GBM*(47)*, we also evaluated the combined effect of DOXY and RT. DOXY monotherapy significantly prolonged survival compared with control (p = 0.008), and the combination of DOXY with RT further extended survival relative to RT alone (p = 0.045) (Fig. 7D). Although survival tended to be longer in the DOXY plus RT group than in the DOXY-alone group, this difference did not reach statistical significance (p= 0.15). No apparent systemic toxicity was observed, and all treatment groups maintained body weights within ±10% of baseline (Fig. 7E). Immunostaining of xenografts showed reduced MT-CO1 expression and increased cleaved caspase-3 levels in DOXY-treated GBMs (Fig. 7F-I), consistent with our findings from in vitro mechanism of action studies. Thus, even at human-equivalent dosing regimens, DOXY exhibited monotherapy antitumor activity, which was further enhanced when combined with RT.

Together, our findings indicate that DOXY impairs mitochondrial translation, reduces mitochondrial membrane potential, and activates OMA1. Activated OMA1 promotes OPA1 proteolysis, leading to mitochondrial fragmentation and ROS accumulation, while simultaneously triggering DELE1 cleavage. Both ROS and DELE1 converge on HRI activation, resulting in phosphorylation of eIF2α and subsequent induction of the ATF4-CHOP-TRIB3 apoptotic pathway (Fig. 7J). This mechanism is selective for GSCs relative to DGCs or non-malignant cells.

Collectively, these findings identify DOXY as a promising therapeutic candidate for GBM. The DOXY dosage used in mice closely corresponds to clinically achievable concentrations in humans, underscoring the translational relevance of our findings. Moreover, combining DOXY with mitophagy inhibition may further enhance its therapeutic efficacy, representing a potential strategy for future investigation. Taken together, these preclinical data highlight the feasibility and near-term translational potential of repurposing DOXY as an adjuvant therapy for GBM.

## DISCUSSION

In this study, we identify a mitochondrial stress response vulnerability that is preferentially engaged in GSCs, but not in DGCs or non-malignant neural cells. We show that this GSC-specific dependence on mitochondrial function renders stem-like tumor cells uniquely susceptible to pharmacological disruption of mitochondrial translation. Our data indicate that treatment-resistant GSCs depend heavily on mitochondrial metabolism for their survival, nominating mitochondrial activity as an appealing therapeutic target. Even within the same genetic background, differentiation reduced mitochondrial dependence. DOXY selectively targeted GSCs and suppressed their proliferation and sphere formation at low doses. Mechanistically, DOXY induced mitochondrial dysfunction, leading to a reduction in mitochondrial membrane potential and activation of the OMA1-DELE1 signaling axis, accompanied by increased production of ROS. These mitochondrial stress signals converged on activation of the HRI-p-eIF2α-ATF4-CHOP pathway, ultimately triggering apoptosis. Finally, the antitumor efficacy of DOXY was validated in patient-derived organoids and in vivo xenograft models at clinically relevant doses, and demonstrated robust synergy with concurrent radiation, a mainstay of the clinical treatment paradigm for GBM. These findings establish mitochondrial translation as a selective vulnerability in GSCs and support the repurposing of DOXY as a clinically applicable therapeutic candidate.

Across transcriptomic, proteomic, and functional analyses, GSCs exhibited evidence of higher mitochondrial activity than DGCs. These data underscore a functionally essential reliance on oxidative metabolism rather than a mere correlative increase. Pharmacological inhibition of OXPHOS markedly reduced GSC viability, and analyses of gene expression patterns in other tumor types (ovarian CSCs) suggest that the mitochondrial dependency of cancer stem-like cells may be independent of the tissue of origin. These findings are consistent with previous reports demonstrating that mitochondrial metabolism underlies the stemness and therapeutic resistance of GSCs*(48)*.

The goal of our research is to develop safe and effective therapeutic strategies aimed at eradicating GSC populations in GBM. Therefore, our drug selection strategy focused on compounds that are BBB penetrant, well tolerated, and preferentially cytotoxic to GSCs. Considering the strong mitochondrial dependency of GSCs we observed, we focused on agents that impair mitochondrial function. Specifically, we selected clinically available antibiotics from distinct structural classes, including tetracyclines, amphenicols, and oxazolidinones*(49)*. Furthermore, we prioritized drugs already approved for central nervous system infections such as meningitis, because these agents have established safety profiles, proven BBB permeability, and are readily accessible for potential clinical repurposing*(50, 51)*. Among the tested compounds, DOXY exhibited a striking and selective cytotoxic effect against GSCs, while showing minimal impact on non-malignant cells. Notably, prolonged exposure to DOXY induced significant GSC death, and concentrations as low as 5 μM suppressed sphere formation over seven days. These results demonstrate that DOXY exerts potent, selective anti-GSC activity at clinically achievable concentrations. Importantly, the clinical familiarity and safety of DOXY make it an ideal candidate for therapeutic repurposing in GBM. Higher DOXY concentrations used in short-term in vitro experiments were employed exclusively for mechanistic dissection and are not intended to model steady-state clinical exposure. Nevertheless, prolonged low-dose exposure within clinically observed cerebrospinal fluid ranges was sufficient to suppress GSC self-renewal.

Mechanistically, DOXY impairs mitochondrial translation, lowers membrane potential, and shifts OPA1 processing toward the short form, promoting mitochondrial fragmentation. Molecular oxygen is the obligate electron acceptor that supports electron transport chain activity. This activity inherently produces superoxide as a byproduct*(52)*. When electron transport is disrupted, fragmented and dysfunctional mitochondria exhibit enhanced electron leakage from the respiratory chain, resulting in excessive superoxide generation*(53)*. These events account for the ROS-dependent component of DOXY-induced apoptosis, as evidenced by the partial rescue of cell death caused by DOXY with antioxidants.

Previous reports have shown that DOXY induces apoptosis through ER stress and activation of the ATF4-PUMA axis in stem-like prostate cancer cells*(17)*. In contrast, oligomycin has been reported to induce mitochondrial dysfunction and activate the OMA1-DELE1-HRI-p-eIF2α-ATF4 signaling cascade in normal cells*(18)*. Unlike these previous studies, we found that in GSCs the ISR response elicited by DOXY occurred independently of PERK. Similar to oligomycin, DOXY caused mitochondrial dysfunction and upregulated the ISR pathway, including p-eIF2α and ATF4, through HRI activation specifically in GSCs, whereas this response was absent in DGCs. The selective activation of the HRI-ATF4 axis in GSCs might reflect their intrinsically high mitochondrial activity, which may predispose them to ISR activation upon mitochondrial perturbation. We delineated this mechanism, encompassing the upstream activation of ATF4 and downstream CHOP-TRIB3 cascade, that links DOXY-induced mitochondrial dysfunction to the ATF4-CHOP-TRIB3 apoptotic pathway. Taken together, our study demonstrates that DOXY-induced apoptosis is mediated through the mitochondria-originated OMA1-DELE1-HRI-p-eIF2α-ATF4-CHOP-TRIB3 pathway rather than the canonical ER stress-PERK-ATF4 axis, uncovering a previously unrecognized vulnerability in GSCs.

Our findings suggest that both autophagy and mitophagy act as protective mechanisms under DOXY-induced mitochondrial stress. Accordingly, combining DOXY with inhibitors of autophagy or mitophagy was synergistically cytotoxic to GSCs. Although bafilomycin is suitable for mechanistic studies in vitro, its toxicity precludes clinical application*(54)*. CQ, an antimalarial agent widely used in humans, was selected as a clinically applicable autophagy inhibitor*(54)*. Previous studies have reported that CQ prolongs survival in GBM patients and clinical trials have been conducted using CQ-based regimens*(55)*. Therefore, further investigation of DOXY and CQ combination therapy in preclinical GBM models is warranted.

To validate the translational relevance of our findings, we confirmed that DOXY exerted consistent antitumor effects in both organoid and in vivo models. The organoid model provided insight into the effects of DOXY within the tumor microenvironment*(56)*. In vivo, a previous study using a subcutaneous DGC mouse model reported that DOXY at 100 mg/kg was effective when combined with TMZ*(19)*. In humans, the clinically used dose of DOXY is 200-300 mg/day*(57)*. According to the NICE guidelines (NG95), DOXY 200 mg twice daily or 400 mg once daily for 21 days is recommended for Lyme disease with CNS involvement*(58)*, which corresponds to approximately 40-80 mg/kg/day in mice based on dose conversion. In our study, we administered DOXY at 40 mg/kg/day. Even at this relatively low dose, DOXY significantly prolonged survival. Reported CSF penetration ratios for DOXY range from 11 to 56%*(29)*, and following administration of 200 mg twice daily in humans, serum concentration reach 3.6-8.6 μg/mL*(29)*. Our data indicate that DOXY potentiates the antitumor efficacy of RT. This result is promising because GSC populations can drive resistance to RT*(9)*. DOXY may mitigate this resistance by specifically targeting GSCs. By confirming efficacy in both patient-derived organoid and orthotopic xenograft models, our study establishes the potential of DOXY-based therapy for GBM. Considering that enhanced mitochondrial activity and ISR signaling are conserved features of multiple cancer stem-like cell populations*(41, 59, 60)*, our findings may have implications beyond GBM and warrant further evaluation in clinically relevant settings. Our findings identify a previously unrecognized mitochondrial stress response vulnerability in GSCs that can be therapeutically exploited using the clinically approved antibiotic DOXY at human-equivalent doses. This work reveals that engagement of a mitochondrion-driven OMA1-DELE1-HRI-p-eIF2α-ATF4-CHOP integrated stress response represents a cell state-specific liability in GSCs with direct therapeutic implications. By establishing the relevance of this pathway across in vitro systems, patient-derived GBM organoids, and orthotopic xenograft models, our results support the translational potential of targeting mitochondrial stress responses to overcome GSC-mediated therapeutic resistance in GBM. These findings suggest that mitochondrial stress adaptation may represent a broadly exploitable vulnerability in stem-like tumor cell population, and an early phase clinical trial at our center is planned to further evaluate these findings in a population undergoing resection of GBM.

### Limitations

Although we calculated equivalent doses for clinical translation, the pharmacokinetics in mice differ from those in humans. Further pharmacokinetic and pharmacodynamic studies, including oral dosing regimens and BBB penetration dynamics, will be essential to fully establish the translational applicability of DOXY. While our study demonstrates the efficacy of DOXY in preclinical models, additional investigations are needed to optimize dosing strategies and evaluate long-term safety in combination regimens. Future studies integrating exposure-response modeling, CNS drug distribution analysis, and trial to evaluate in vivo effects and define optimal therapeutic windows.

## MATERIALS AND METHODS

### Experimental design

Sample sizes were based on prior experience and published studies. Biological replicates represent independent experiments or animals. No animals or organoids were excluded. Randomization was performed prior to tumor implantation; blinding was not feasible for in vivo studies. Survival endpoints were defined a priori. For immunohistochemical analyses, tumors of comparable size and anatomical location were quantified.

### Human subjects

Human glioma tissue was obtained with informed consent under protocols approved by the University of Pittsburgh Medical Center Institutional Review Board, in accordance with the Declaration of Helsinki. Samples were de-identified prior to analysis and classified according to the 2021 World Health Organization (WHO) Classification of Tumors of the Central Nervous System*(61)*. Patient-derived tumor explants and organoids were generated as previously described *(45)*.

### Cell lines and culture conditions

Patient-derived GSCs and DGCs were obtained from established sources and maintained under standard conditions as previously reported *(62, 63)*. Normal human astrocytes (NHAs) and non-glioma cell lines were cultured according to the suppliers’ recommendations *(15)*. Cell line sources, sex (when available), and detailed culture conditions are provided in the Supplementary Methods. Cells were routinely tested for mycoplasma contamination.

### Animals

All animal studies were approved by the University of Pittsburgh Medical Center Institutional Animal Care and Use Committee and conducted in accordance with institutional and federal guidelines. ICR-SCID mice were housed under pathogen-free conditions with ad libitum access to food and water.

### Orthotopic xenograft models and treatments

Orthotopic glioma xenografts were established by intracranial implantation of patient-derived GSC387 cells into adult ICR-SCID mice. Mice were assigned to treatment groups receiving saline, doxycycline (DOXY), radiation therapy (RT), or combination therapy. DOXY was administered at a dose of 40 mg/kg/day, initially by intraperitoneal injection and subsequently via oral administration. Fractionated RT (2 Gy/day) was delivered on consecutive days as indicated. Animals were monitored daily and euthanized upon development of neurological symptoms. Detailed surgical procedures and treatment schedules are described in the Supplementary Methods.

### Pharmacological treatments

DOXY and additional pharmacological agents were selected based on blood-brain barrier permeability, clinical safety profiles, and relevance to mitochondrial function. Complete lists of reagents, antibodies, and sources are provided in Supplementary Table 1.

### Patient-derived organoids

Glioblastoma organoids were generated from freshly resected patient tumor tissue and maintained under defined culture conditions to preserve tumor architecture and cellular heterogeneity *(45)*. Organoids were treated with DOXY and analyzed for viability, apoptosis, and molecular signaling responses as described in the Supplementary Methods.

### Statistical analysis

Statistical analyses were performed using GraphPad Prism and R. Data are presented as mean ± SD unless otherwise indicated. Comparisons between two groups were performed using unpaired two-tailed Student’s t-tests or non-parametric tests as appropriate. Comparisons among multiple groups were conducted using one-way ANOVA with appropriate post hoc corrections. Survival data were analyzed using Kaplan-Meier curves and log-rank tests. P values <0.05 were considered statistically significant. Additional details regarding statistical analyses and replicate definitions are provided in the figure legends and Supplementary Methods.

## Acknowledgements

This study was supported by National Institutes of Health (NIH) grants R01CA258586 and R01CA289260 to S.K.M. and K.G.A., R01GM158820 and R01NS142141 to S.K.M., P50CA165962 and U19CA264504 to S.K.M. This work was also supported by awards from Cancer Prevention and Research Institute of Texas (CPRIT) grants RP240489 and RP230344 to S.K.M., Japan Society for the Promotion of Science (JSPS) Overseas Research Fellowship to K.M., the Uehara Memorial Foundation to K.M., the Mochida Memorial Foundation for Medical and Pharmaceutical Research to K.M., the Alumni Association of the Faculty of Medicine, Kyushu University Overseas Research Fellowship to K.M.

## Author contributions

Conceptualization: KM, SKM, KGA

Methodology: KM, MTG, FY, FL, NM, SO, KY, POZ, SA, SKM, KGA

Investigation: KM, MTG, FY, FL, NM, SO, KY, POZ, SA, SKM, KGA

Formal analysis: KM

Data curation: KM

Software: KM

Supervision: KY, POZ, SA, SKM, KGA

Project administration: SKM, KGA

Writing-original draft: KM, MTG, SKM, KGA

Writing-review and editing: KM, MTG, SKM, KGA

Funding acquisition: SKM, KGA

## Data and materials availability

Publicly available RNA-seq datasets analyzed in this study are available through GEO under accession number GSE54791. Newly generated RNA-seq data will be deposited in GEO and made available upon publication.

## Supplementary Information

### Methods

#### Human Subjects

The study was conducted according to the principles of the Declaration of Helsinki. Patient tissue was collected following ethical and technical guidelines on the use of human samples for biomedical research at the University of Pittsburgh Medical Center after informed patient consent under a protocol approved by the University of Pittsburgh Medical Center’s Institutional Review Board. All patient samples were de-identified before processing. All patient samples and organoids were diagnosed and graded according to the 2021 WHO Classification of Tumors of the Central Nervous System (CNS), 5th edition*(1)*. Tissue explants from tumor resection surgeries were generated as previously described*(2)* and including patients with the following ages, sexes and clinical diagnoses: UPMC917 (72-year-old male; Glioblastoma, IDH-wildtype). Higher DOXY concentrations used in selected in vitro experiments were employed exclusively for mechanistic interrogation and are not intended to model clinically achievable steady-state exposure.

#### Cell lines

NHA cells were generated from commercially available astrocytes (Lonza, CC-3187) as described previously*(3)*. HEK293T cells (female, ATCC CRL-3216), U251, A172, U138 were obtained commercially. TS516 (sex not reported, RRID: CVCL_A5HY) was provided by I. Mellinghoff at Memorial Sloan-Kettering Cancer Center*(4)*. GSC23, GSC387, GSC3028, GSC3565, and NSC11 (sex not reported) cells were provided by J. Rich at University of North Carolina-Chapel Hill*(5)*. GSC63 (UTSW63) (male) was generated as previously described*(6)*. GSC5 (UTSW5) was generated from a female glioblastoma patient at UT Southwestern*(6)*. All cell lines were routinely evaluated for mycoplasma contamination with the e-Myco Mycoplasma PCR Detection Kit (Bulldog Bio, 2523348), e-Myco PLUS Mycoplasma PCR Detection Kit (Bulldog Bio, 25233), or MycoAlert Mycoplasma Detection Kit (Lonza, LT07-318), per manufacturer’s instructions. As reference short term tandem repeat profiles have not been established for these lines, no cell line authentication was performed. Sex and source of each line are stated above or listed as unknown if unreported in the original publication describing its derivation.

#### Cell Culture

GSC23, GSC387, GSC3028, GSC3565, TS516, GSC5 (UTSW5), and GSC63 (UTSW63) human GSCs were cultured in NeuroCult NS-A Basal Medium (Human) with 1x Proliferation Supplement (STEMCELL Technologies 05751), supplemented with 20 ng/mL EGF (STEMCELL Technologies, 78006); 20 ng/mL bFGF (STEMCELL Technologies, 78003); 2 µg/mL heparin (STEMCELL Technologies, 07980); 100 U/mL and 100 μg/mL, respectively, of penicillin/streptomycin (Thermo Fisher Scientific, 15140-122); 250 ng/mL amphotericin B (Cytiva SV30078.01); and 0.25 µg/mL Plasmocin (InvivoGen, MPP-43-04) on ultra-low adherence plates (6-well plates: Corning, 3471, 10 cm dishes: Corning, 4615) in 5% CO2 and at ambient oxygen at 37°C. To minimize batch-to-batch variability, experiments directly comparing treatment conditions were performed using the same lot of NeuroCult medium whenever possible. NSC11 cells were cultured in Neurobasal Medium-A (Thermo Fisher Scientific, 10888-022) supplemented with GlutaMAX (Thermo Fisher Scientific, 35050-061); sodium pyruvate (Thermo Fisher Scientific, 11360-070) 1x B27 supplement (Thermo Fisher Scientific, 12587001); 20 ng/mL EGF; 20 ng/mL bFGF; 2 µg/mL heparin; 50 U/mL and 50 μg/mL, respectively, of penicillin/streptomycin; on ultra-low adherence plates (6-well plates: Corning, 3471, 10 cm dishes: Corning, 4615) in 5% CO2 and at ambient oxygen at 37°C.

NHA, A172, U138, and U251 were cultured adherently in DMEM (Thermo Fisher Scientific, 11995-065) supplemented with 10% fetal bovine serum (FBS; Thermo Fisher Scientific, 12999102) and 100 U/mL and 100 μg/mL, respectively, of penicillin/streptomycin (Thermo Fisher Scientific, 15-140-148) in 5% CO2 and at ambient oxygen at 37°C.

#### Animals

All care and treatment of experimental animals were carried out in strict accordance with Good Animal Practice as defined by the US Office of Laboratory Animal Welfare and approved by the University of Pittsburgh Medical Center Institutional Animal Care and Use Committee (protocol 25066832). Animal welfare assessments were carried out daily during treatment periods. Animals were housed in a pathogen-free environment between 20-26°C and at 30-70% humidity, with a 12-hour:12-hour light: dark cycle. ICR-SCID (Taconic) mice were obtained from Taconic at 6-7 weeks of age. Mice were housed together (2-5 mice of the same sex per cage) and provided free access to water and chow diet (Purina ISO Pro Rodent 3000).

#### Generation of patient-derived xenograft (PDX) mouse models

Female mice were housed together (2-5 mice per cage) and provided free access to standard diet and water. GSC387 orthotopic xenografts were established by intracranial injection of 3,000 cells into 10-week-old female ICR-SCID mice (IcrTac:ICR-*Prkdc^scid^*, Taconic ICRSC). For orthotopic glioma cell implantations, mice were anesthetized via isoflurane and immobilized using a stereotactic frame. An incision was made to expose the skull surface, and a hole was drilled into the skull. Cells suspended in 5 µL HBSS (Thermo Fisher Scientific, 14175-095) were injected into the brain through the hole using a 5 μL syringe (Hamilton). 0.5-1 mm anterior and 2 mm lateral to the bregma, at a depth of 3 mm from the brain surface. The skin was closed with surgical clips, and buprenorphine was given for analgesia. Survival analyses were performed by researchers who were not blinded to the treatment arms or genotypes of the mice. Mice were euthanized upon displaying neurological symptoms or moribundity. Mice were assigned to one of the following treatment groups: saline, DOXY (40 mg/kg/day), radiation (2 Gy×3), DOXY + radiation. Starting the day after tumor cell injection, DOXY (40 mg/kg/day) was administered by IP injection five days per week. Eight days after injections, the regimen was switched to the oral administration. For oral dosing, DOXY was dissolved in autoclaved drinking water at a final concentration of 200 mg/L, filter-sterilized and provided ad libitum in light-protected bottles (foil-wrapped). Solutions were prepared fresh and replaced daily. Assuming a 25g body weight and typical cage water consumption, this regimen yields an estimated intake of 1 mg/mouse/day (40 mg/kg/day). For the RT group, mice received fractionated irradiation (2Gy/day) on days 7-9 after injection. Animals were monitored daily until the appearance of neurological symptoms, at which point they were euthanized.

#### Chemicals

All reagents and antibodies used are shown in Supplementary Table 1.

#### Plasmids and cloning

shRNAs utilized in this research include: shATF4: sh1TRCN0000013573, sh2TRCN0000013575; shCHOP: sh1TRCN0000007263, sh2TRCN0000007264; shTRIB3: sh1TRCN0000037404, sh2TRCN0000037408; shHRI: sh1TRCN0000010231, sh2TRCN0000196718; shOMA1: sh1TRCN0000062118, sh2TRCN0000062120. A non-targeting shRNA that does not target any known human transcript (Sigma-Aldrich, SCH002, shCONT) was the negative shRNA control.

#### Retroviral packaging and infection

For the stable expression of selected targets in GSCs, we conducted virus packaging and infection using recombinant lentivirus. A mixture of transfer plasmid, psPAX2 (Addgene), pMD2.G (Addgene), and OPTI-MEM (Thermo Fisher Scientific, 31985-062) was co-transfected using Lipofectamine 2000 (Invitrogen, 11668027) into HEK293T cells. Supernatants were collected at 48 and 72 h post-transfection, concentrated using Lenti-X concentrator (Takara, 631232). GSCs were infected with virus + 10 μg/ml polybrene (Sigma, H9268) for 24 h and selected with 2 μg/ml puromycin for 2 days. The efficiency of infection was assessed through qPCR or immunoblotting.

#### Immunoblot analysis of protein expression

Cells were lysed in RIPA buffer (Thermo Fisher Scientific, 89900) containing PMSF (CST, 8553) and PhosSTOP (Roche, 04906845001). Lysates were centrifuged (15,000 rpm, 10 min, 4℃), quantified by BCA assay (Thermo Fisher Scientific, 23227), and resolved by SDS-PAGE. Proteins were transferred to nitrocellulose membranes (Bio-Rad, 1620112) and blocked with Blocking One (Nacalai Tesque, 03953-95). Primary and secondary antibodies were used per manufacturer’s protocols, signals detected with SuperSignal West Femto (Thermo Fisher Scientific, 34095) or Immobilon ECL Ultra (Sigma, WBULS0100) on a ChemiDoc or iBright 1500.

#### Quantitative real-time PCR

RNA was isolated using RNeasy Mini Kit (QIAGEN, 74104) and reverse-transcribed with iScript cDNA Synthesis Kit (Bio-Rad, 1708891). qPCR was performed with iTaq Universal SYBR Green Supermix (Bio-Rad, 1725120) on a CFX96 system. 18S rRNA or β-actin served as control. Primer sequences are listed in Supplementary Table 2.

#### Seahorse XF96 flux analyzer

Mitochondrial OXPHOS and glycolytic activity were measured using the Seahorse XF96 Analyzer (Agilent Technologies). Seahorse XF96 microplates were pre-coated with Matrigel (Corning, 354277) for 30 min at 37℃ before cell seeding. GSCs (2 × 10^4^ cells/well) and DGCs (1× 10^4^ cells/well) were plated and incubated for approximately 24 h prior to analysis. Basal OCR and ECAR were measured according to the manufacturer’s protocol. Sequential injections of oligomycin (1.5 μM), FCCP (0.5 μM+1.0 μM), and rotenone and antimycin (0.5 μM) were performed to assess mitochondrial bioenergetic parameters. Following the assay, cells were trypsinized and counted using the Countess Automated Cell Counter (Invitrogen), or quantified based on DNA content using the CyQUANT Cell Proliferation Assay Kit (Thermo Fisher Scientific, C7026). Data were normalized to cell number determined immediately after the assay.

#### Flow cytometry (TMRE, MitoSOX, CellROX)

Cells were collected, gently dissociated, and resuspended in HBSS. For mitochondrial membrane potential, cells were incubated with TMRE (25 nM, 20 min, 37℃, protected from light), washed once with HBSS. FCCP was used as a positive control. For mitochondrial superoxide detection, cells were incubated with MitoSOX Red (2.5 μM, 10 min, 37℃), washed with HBSS, and analyzed immediately. For cellular ROS, cells were incubated with CellROX Deep Red (2.5 μM, 20 min, 37℃), washed with HBSS, and analyzed immediately. Data were acquired using a Beckman Coulter CytoFLEX 6L flow cytometer with the following detection channels: TMRE (PE), MitoSOX (mCherry/Y610), CellROX (APC). Forward and side scatter (FSC/SSC) were used to gate single, viable cells and exclude debris. Unstained controls were used to set negative populations. Because DGC387 cells exhibited substantially larger cell bodies than GSC387 cells, TMRE intensity for the GSC387-DGC387 pair was normalized to the estimated cell volume as inferred from FSC to correct for size-dependent differences in integrated fluorescence.

#### Apoptosis assay

Apoptosis was assessed using FITC AnnexinV (BD Biosciences, 51-65874) and PI (Invitrogen, R37610) according to the manufacturer’s instructions. Samples were analyzed on a Beckman Coulter CytoFLEX 6L flow cytometer. FSC/SSC were used to gate cells and exclude debris. Negative samples were used to define the negative population.

#### mt-Keima experiment

Mitophagy was assessed using the pH-sensitive fluorescent protein mt-Keima, as previously described*(7)*. The pHAGE-mt-Keima plasmid (Addgene plasmid, #131626) was used for stable expression in GSC387 cells. Cells were transfected with the plasmid, and after confirming mt-Keima fluorescence by microscopy, they were used immediately for subsequent assays. For live-cell imaging, mt-Keima fluorescence was visualized using a confocal microscope. Under neutral conditions, mt-Keima is excited at 440 nm and emits at 620 nm, whereas under acidic conditions it is excited at 560 nm and emits at 620 nm. For flow cytometry, mt-Keima fluorescence was analyzed in the Y610 (mCherry) channel. Mitophagy-associated events were defined based on a shift toward higher mt-Keima signal, with gates set using untreated cells to define the negative population and FCCP-treated cells as a reference for mitochondrial stress-associated shifts on a Beckman Coulter CytoFLEX 6L flow cytometer.

#### Immunostaining

Cells (2 × 10^4^ per dish) were seeded on glass bottom dishes and cultured. Cells were fixed with 4% paraformaldehyde phosphate buffer solution for 10 min, permeabilized with 0.1% Triton X-100/PBS for 5 min, and blocked with Blocking One (Nacalai Tesque). Primary antibodies were diluted with Can Get Signal (Toyobo) and incubated for 1 h at room temperature, followed by secondary antibody incubation (Invitrogen: anti-mouse IgG Alexa Fluor 594 (A11032), anti-rabbit IgG Alexa Fluor 594 (A11012), and anti-rabbit IgG Alexa Fluor 488 (A11008). Slides were mounted with ProLong Diamond Antifade Mountant (Thermo Fisher Scientific, P36965) and imaged under a fluorescence microscope (Nikon Eclipse Ti2 or Nikon A1R).

#### Organoid creation

Organoids from GBM patients were generated as previously reported*(2)*. Briefly, tumor tissue was collected from the operating room, directly suspended in ice-cold Hibernate A and transferred to the laboratory on ice within 30 minutes of explantation. Tumor pieces were moved into RBC lysis buffer (Thermo Fisher 00433357) and incubated at room temperature for 10 minutes with rocking. Tumor pieces were then washed with Hibernate A containing GlutaMAX, penicillin/streptomycin, and amphotericin B. Tissues were cut and plated, one per well, in a 24-well ultra-low adherence plate (Corning 3473) in 1 mL Glioma Organoid Complete Medium (GOC)*(8)*.

#### Histopathology of human explants

Human SXOs were fixed for 1 h in 10% neutral buffered formalin (Sigma, HT501128). After fixation, tissues were washed, stored in 70% ethanol, paraffin embedded, sectioned, and stained with hematoxylin and eosin (H&E) by the UPMC Histopathology Core.

#### Immunohistochemistry Staining

Paraffin-embedded tissues were sectioned at 5 μm thickness. Antigen retrieval was performed in citrate buffer, followed by blocking with Blocking One (Nacalai Tesque) for 30 min. Sections were incubated overnight at 4℃ with the following primary antibodies: MT-CO1 (1:5000, abcam, #14705), SOX2 (1:300, CST, #14962) and cleaved caspase-3 (1:200, CST, #9661). Signals were visualized using a DAB substrate kit (CST, 8059). Images were acquired using a Nikon Eclipse Ti2 microscope, and quantification was performed using ImageJ by calculating the intensity from random fields per sample.

#### Cell growth assay

Cells were plated in 96-well plates as follows: for 24-72 h assays: GSCs, 1.5 x 10^4^ cells/well; DGCs, 5 x 10^3^ cells/well, for 96 h assays: GSCs: 5 x 10^3^ cells/well; DGCs, 2 x 10^3^ cells/well. Cells were cultured in NeuroCult NS-A Basal Medium (Human) or DMEM as described above. Cell viability was measured at 24, 48, 72, and 96 h using CellTiterGlo (Promega, G7570). Media in the remaining wells were replaced at 48 h. Growth rates were analyzed by nonlinear regression to fit exponential growth curves in GraphPad Prism.

#### Public RNA-seq analysis

Publicly available RNA-seq data comparing GSCs and DGCs were obtained from the GEO database (GSE54791). CPM normalized expression matrices were analyzed in R (v4.3.2). For GSEA, genes were ranked based on Welch’s t-statistics comparing GSCs versus DGCs, and enrichment analysis was performed using fgsea package with 10,000 permutations. Hallmark and Reactome gene sets, including OXPHOS, TCA cycle, and RET, were obtained from MSigDB via the msigdbr package. For GO enrichment analysis, differential expression was assessed using the limma package following log2 transformation of CPM values. Genes significantly upregulated in GSCs (log2 fold-change>log2(1.5), adjusted P<0.05) were subjected to GO BP enrichment using clusterProfiler::enrichGO, with all expressed genes used as the background universe. From the enriched GO terms, those related to mitochondrial processes were extracted by filtering for terms containing the keyword ‘mitochondr’ and visualized as dot plots. To extend these findings to another cancer type, publicly available ovarian cancer RNA-seq raw count data from CSCs and non-CSCs were obtained from GSE148003. Differential gene expression analysis was performed using edgeR after filtering genes with CPM>1 in at least two samples. Following normalization and dispersion estimation, differential expression was assessed using a generalized linear model and likelihood ratio test. Genes upregulated in CSCs (log2 fold-change>0, FDR<0.1) were subjected to GO BP and Cellular Component enrichment using clusterProfiler, and mitochondria-related GO terms were extracted and visualized*(9)*.

#### RNA sequencing and transcriptional subtype classification of human primary glioma samples

Total RNA from primary human glioma samples was extracted using the RNeasy Plus Universal kit (Qiagen, 73404) according to the manufacturer’s instructions. Libraries were prepared and sequenced as paired-end 150bp reads on an Illumina platform. Raw FASTQ files were adapter and quality trimmed using Trim Galore, then aligned to the human reference genome (GRCh38/hg38) using HISAT2 with default parameters. Gene-level counts were generated with featureCounts. Downstream analysis was performed in R (v4.3.2) using DESeq2 for differential expression analysis (DOXY treated vs control, with biological replicates per condition). Low-count genes were filtered using DESeq2’s default prefiltering. Counts were rounded to integers, and size-factor and dispersion estimation were followed by Wald tests using the DESeq function. Differentially expressed genes (DEGs) were defined as those with FDR<0.05 and log2 fold-change>1. DEG lists were subjected to pathway enrichment analyses (GO Biological Process, KEGG, and Reactome) using clusterProfiler and ReactomePA. GSEA was performed using the fgsea package based on gene rankings defined by log2 fold-changes. ER stress-related signaling was assessed using the GO:0034976 gene set, and autophagy related signaling was examined using the GO:0006914 gene set obtained from MSigDB. All enrichment results were visualized using ggplot2 and enrichplot. Pathway activity was evaluated using rank-based gene set enrichment analysis rather than by direct averaging of expression values across pathway genes.

#### Patient database bioinformatics

The GlioVis data portal (https://gliovis.bioinfo.cnio.es/) was used to interrogate the clinical relevance of patients with glioma from TCGA data sets. The survival of patients in each group was analyzed using the Kaplan-Meier method via log-rank test.

#### Other statistical analyses

Information related to data presentation and statistical analysis for individual experiments can be found in the corresponding figure legends. Statistical analyses were carried out using GraphPad Prism software. For comparisons between two groups, either parametric (unpaired Student’s t-test) or non-parametric (Mann-Whitney) tests were applied depending on data distribution. For comparisons among three or more groups, one-way ANOVA followed by Sidak’s, Dunnett’s, or Tukey’s multiple comparison tests were used. Survival data from mouse xenograft studies were analyzed and statistical significance was determined using log-rank tests. For all tests, *p-*values < 0.05 were considered statistically significant.

---

**Fig. S1.**
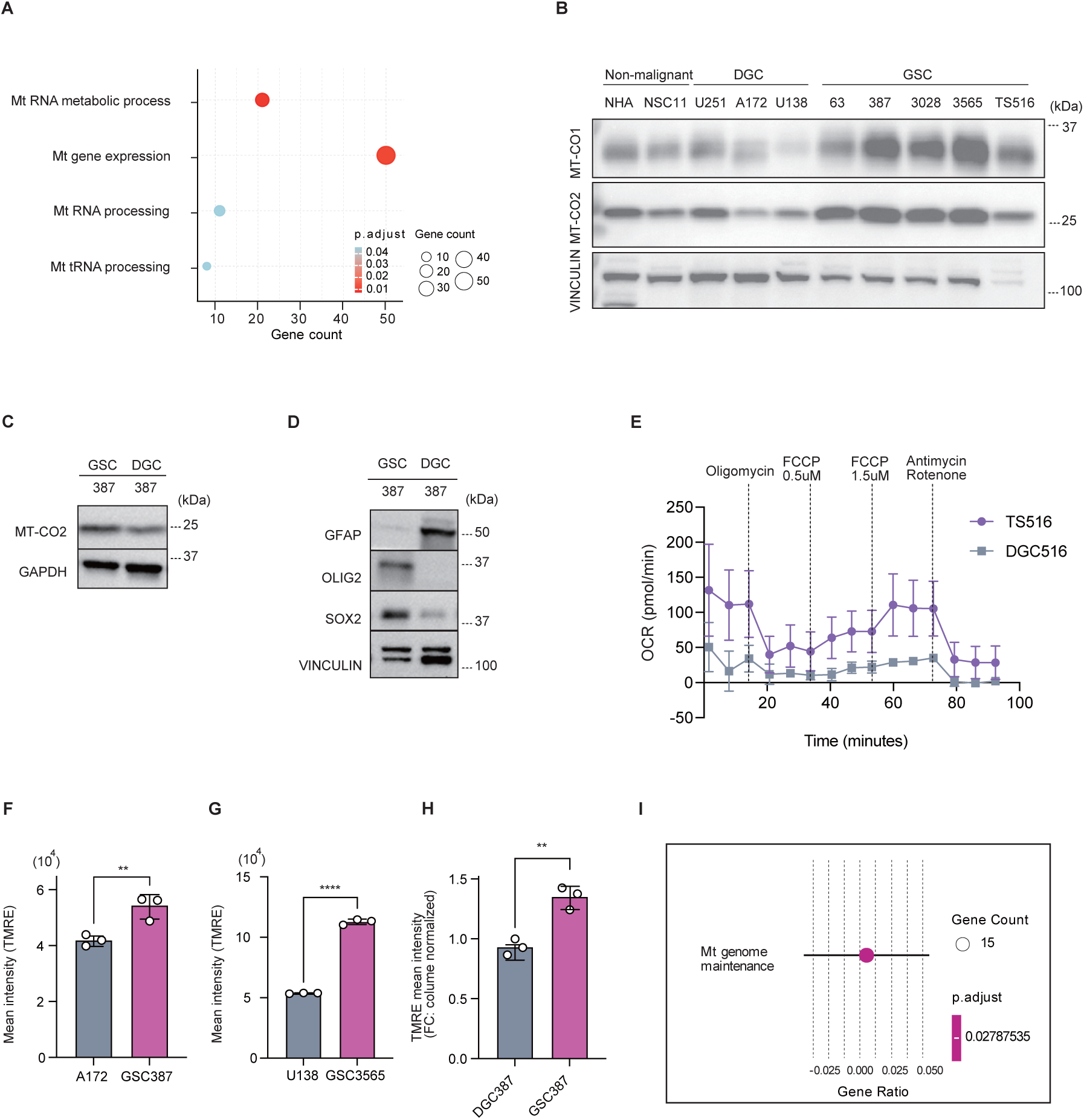
GSCs are dependent on mitochondrial metabolism. (A) GO enrichment analysis of GSCs versus DGCs, highlighting mitochondrial pathways. (B) Western blot analysis of non-malignant cells (NHA, NSC11), DGCs (U251, A172, U138), and GSCs (GSC63, 387, 3028, 3565, TS516), showing expression of MT-CO1, MT-CO2, and VINCULIN (n = 3). (C-D) Western blot analysis of GSC387 and DGC387 showing expression of MT-CO2, GAPDH, GFAP, OLIG2, SOX2, and VINCULIN (n = 3). (E) Oxygen consumption rate (OCR) profiles of GSC516 and DGC516 (n = 5, 6). (F-H) Flow cytometry analysis of mitochondrial membrane potential using TMRE in (F) A172 vs. GSC387, (G) U138 vs. GSC3565, and (H) GSC387 vs. DGC387 (n = 3). (I) GO enrichment analysis of CSCs versus non-CSCs, highlighting mitochondrial pathway in ovarian cancer. Data are presented as mean ± SD. Statistical analyses were performed using an unpaired two-tailed Student’s *t*-test (F-H). **p < 0.01, ****p < 0.0001.

**Fig. S2.**
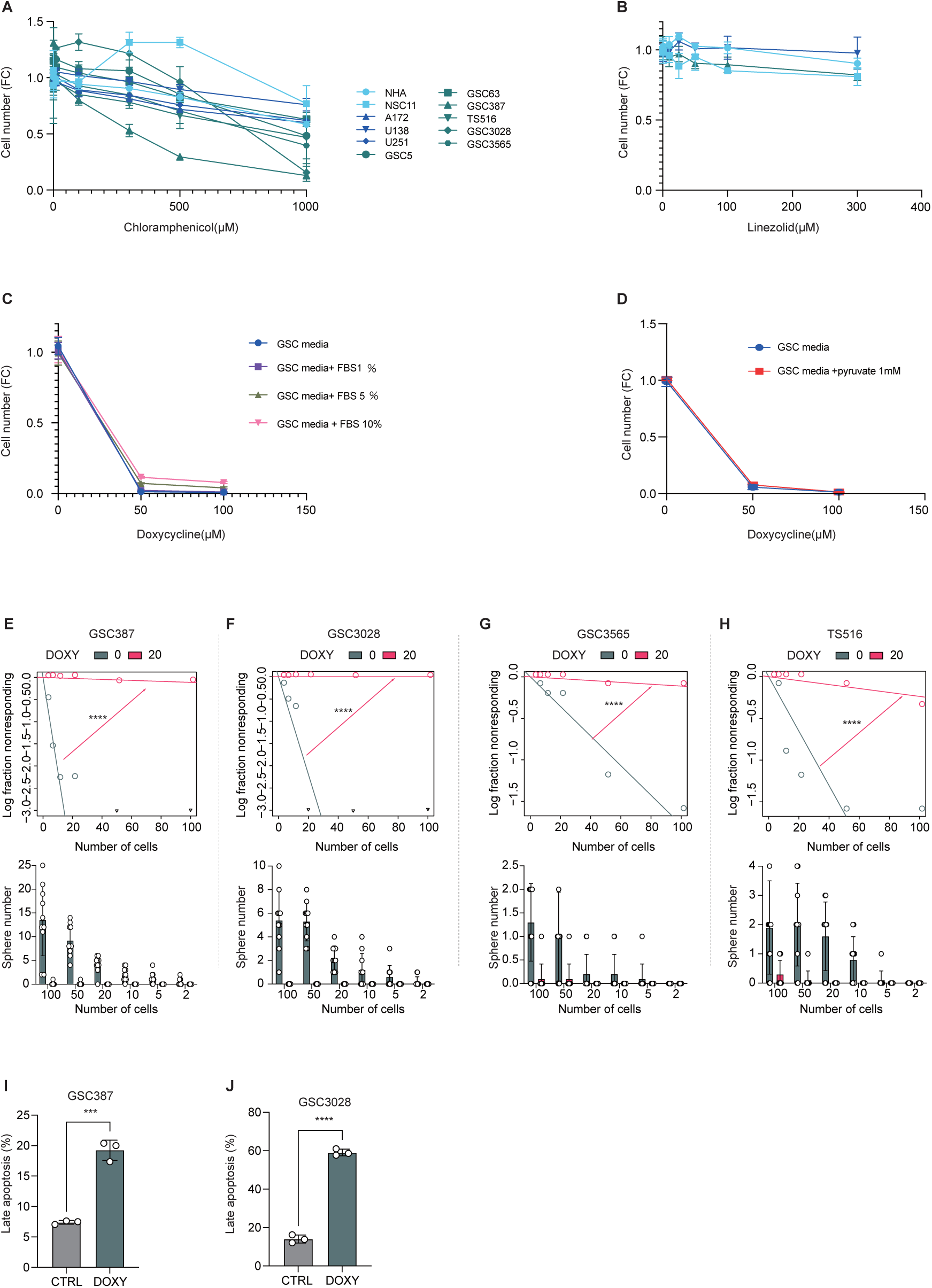
Mitochondrial dysfunction selectively affects GSCs and induces apoptosis. (A) Data represent the same experiment set as Fig. 2C. Cell viability after 72 h of chloramphenicol treatment (n = 3). (B) Cell viability after 48 h of linezolid treatment (n = 3-6). (C) Cell viability after 24 h of DOXY treatment under GSC culture conditions supplemented with increasing concentrations of FBS (0,1,5, and 10%) (n = 4). (D) Cell viability after 24 h of DOXY treatment under GSC culture conditions supplemented with pyruvate (1 mM) (n = 3). (E-H) Extreme limiting dilution assay and sphere-formation assay of GSC387 (E), GSC3028 (F), GSC3565 (G), and TS516 (H) treated with control or 20 μM DOXY for 7 days. (I-J) Flow cytometry analysis of apoptosis in GSC387 (I, 30 μM, 16 h) and GSC3028 (J, 60 μM, 16 h) with or without DOXY treatment (n = 3). Data are presented as mean ± SD. Statistical analyses were performed using an unpaired two-tailed Student’s *t*-test (G, H). ***p < 0.001, ****p < 0.0001.

**Fig. S3.**
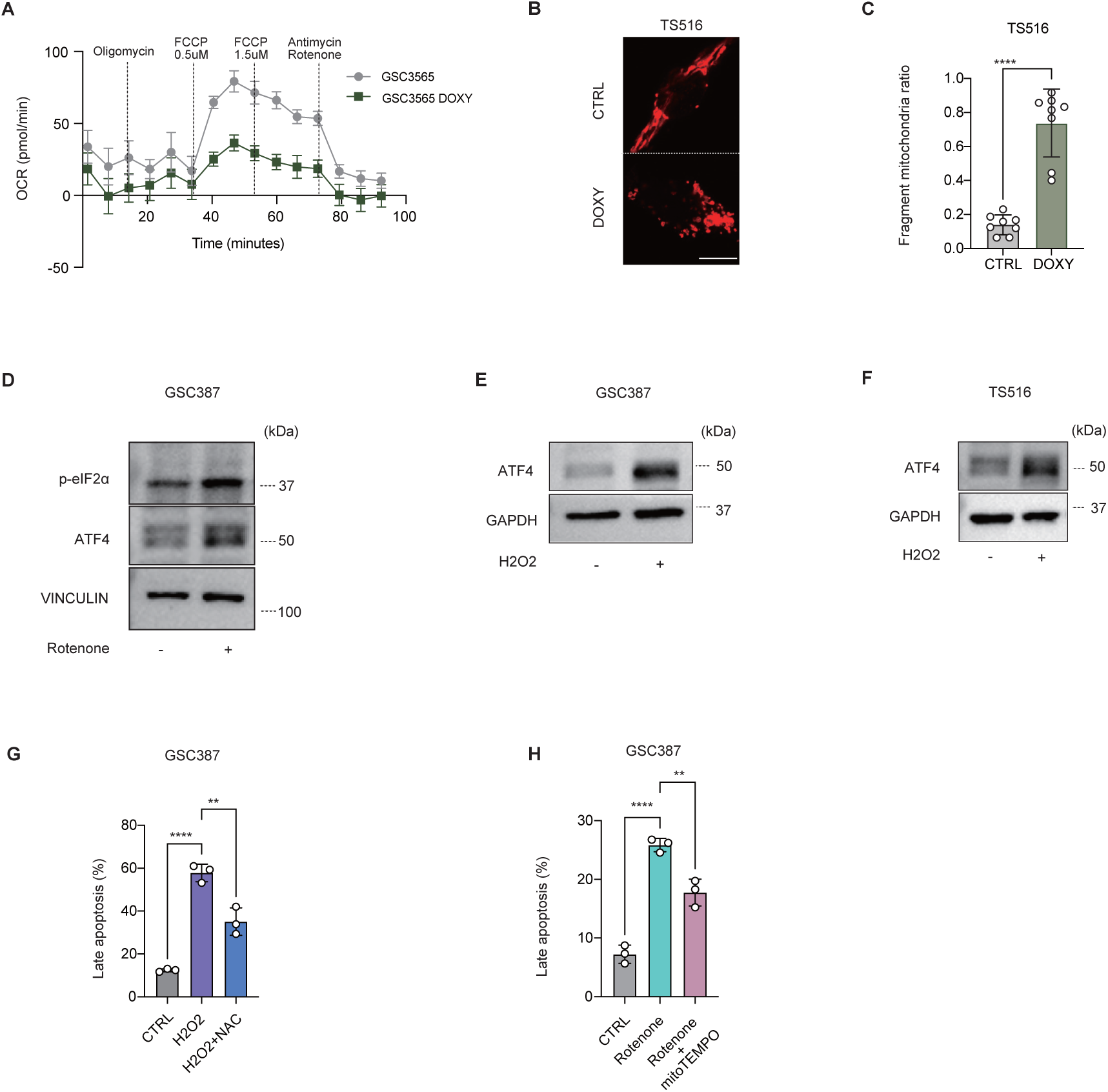
Doxycycline decreases oxygen consumption rate and induces mitochondrial fragmentation. (A) OCR profiles of GSC3565 with or without DOXY treatment (30 μM, 2 h) (n = 4-5). (B) Immunofluorescence images of TOM20 in TS516 cells treated with or without DOXY (30 μM, 16 h). Scale bar: 10 μm. (C) Quantification of mitochondrial fragmentation from (B) (n = 8). (D-F) Western blot analyses of GSC387 treated with or without rotenone (10 μM, 3 h) (n = 3) (D), H₂O₂ (400 μM, 3 h) (E), and TS516 treated with or without H₂O₂ (400 μM, 3 h) (F). (G) Flow cytometry analysis of apoptosis in GSC387 under control, H₂O₂ (400 μM), and H₂O₂ (400 μM) + NAC (0.5 mM) conditions for 4 h (n = 3). (H) Flow cytometry analysis of apoptosis in GSC387 under control, rotenone (10 μM), and rotenone (10 μM) + mitoTEMPO (20 μM) conditions for 24 h. Data are presented as mean ± SD (n = 3). Statistical analyses were performed using an unpaired two-tailed Student’s *t*-test (C) or one-way ANOVA (G, H) followed by Sidak’s multiple comparisons test. **p < 0.01; ****p < 0.0001.

**Fig. S4.**
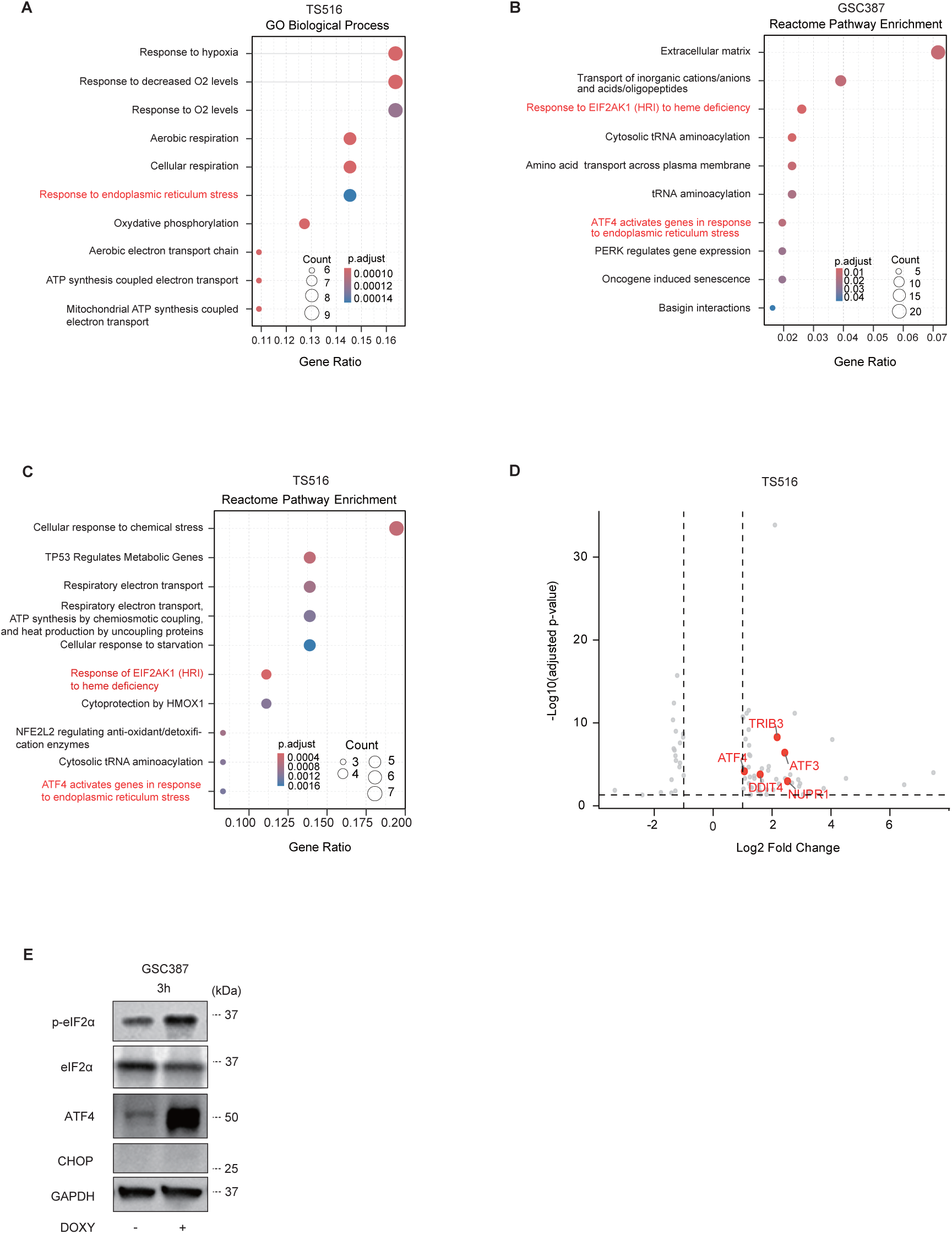
Doxycycline activates the integrated stress response (ISR) in GSCs. (A) GO analysis of RNA-seq data from TS516 treated with or without DOXY (25 μM, 24 h). (B-C) Reactome pathway analysis of RNA-seq data from GSC387 (B) and TS516 (C) treated with or without DOXY. (D) Volcano plot of differentially expressed genes in TS516 with or without DOXY treatment (25 μM, 24 h). (E) Representative western blots showing ATF4-CHOP pathway activation under the indicated conditions. (E) GSC387 with or without DOXY (30 μM, 3 h) (n = 3).

**Fig. S5.**
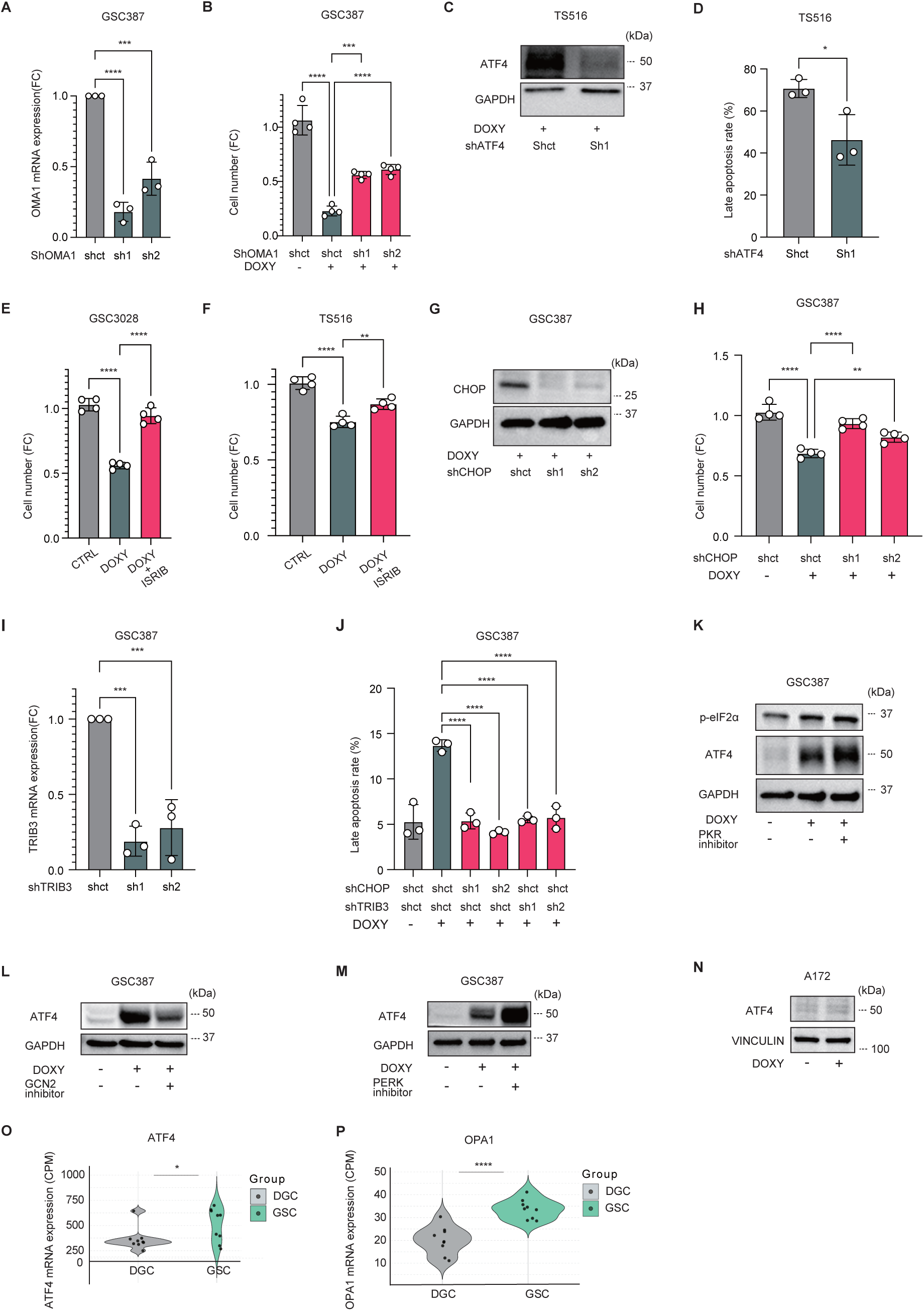
Doxycycline-induced ISR activation triggers apoptosis through the HRI-p-eIF2α-ATF4-CHOP-TRIB3 axis. (A) mRNA expression of OMA1 in shCtrl and shOMA1 cells (n = 3). (B) Cell viability of GSC387 with DOXY (30 μM, 24 h) in shCtrl or shOMA1 cells (n = 4). (C, G, K-N) Representative western blots. (C) TS516 with or without DOXY (50 μM, 3 h) and shATF4. (D) Flow cytometry analysis of apoptosis in TS516 with or without shATF4 (50 μM, 24 h) (n = 3). (E-F) Cell viability of GSC3028 (E) and TS516 (F) treated with control, DOXY (50 μM for GSC3028; 100 μM for TS516), and DOXY + ISRIB (1 μM) for 24 h (n = 4). (G) Western blot analysis of GSC387 with or without DOXY (30 μM, 3 h) and shCHOP. (H) Cell viability of GSC387 with DOXY (30 μM, 24 h) in shCtrl or shCHOP cells (n = 4). (I) mRNA expression of TRIB3 in shCtrl and shTRIB3 cells (n = 3). (J) Flow cytometry analysis of apoptosis in shCtrl, shCHOP, and shTRIB3 cells (30 μM, 16 h) (n = 3). (K) GSC387 treated with or without DOXY (30 μM, 3 h) and PKR inhibitor (1 μM). (L) GSC387 treated with or without DOXY (30 μM, 3 h) and GCN2 inhibitor (2 μM). (M) GSC387 treated with or without DOXY (30 μM, 3 h) and PERK inhibitor (1 μM). (N) A172 treated with or without DOXY (30 μM, 3 h). (T-U) Quantitative comparison of (O) ATF4 and (P) OPA1 expression between DGCs and GSCs based on RNA-seq data. Data are presented as mean ± SD. Statistical analyses were performed using an unpaired two-tailed Student’s *t*-test or one-way ANOVA followed by Sidak’s multiple comparisons test. *p < 0.05; **p < 0.01; ***p < 0.001; ****p < 0.0001.

**Fig. S6.**
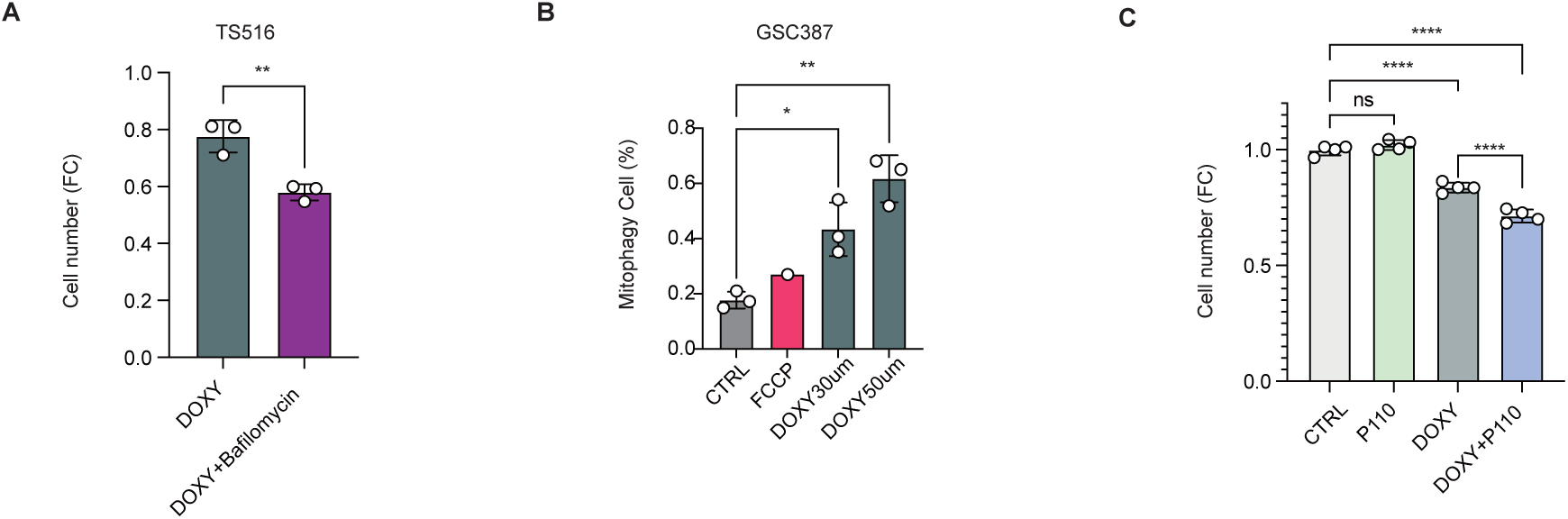
Mitophagy acts as a cytoprotective response to doxycycline induced mitochondrial stress in GSCs. (A) Cell viability of TS516 treated with DOXY (30 μM) with or without bafilomycin (20 nM) for 24 h (n = 3). (B) Flow cytometry analysis of mt-Keima-expressing GSC387 under control, FCCP (10 μM), or DOXY (30 μM, 50 μM) treatment for 16 h (n = 3). (C) Cell viability of GSC387 treated with DOXY (30 μM) with or without p110 (0.5 μM) for 24 h (n = 4). Data are presented as mean ± SD. Statistical analyses were performed using an unpaired two-tailed Student’s *t*-test (A) or one-way ANOVA followed by Sidak’s multiple comparisons test (B, C). *p < 0.05; **p < 0.01; ****p < 0.0001.

**Table S1.** Reagents and antibodies used in this study.

| Reagent or Resource | Source | Identifier |
| --- | --- | --- |
| **Antibodies** |  |  |
| Mouse monoclonal anti-MT-CO1 | abcam | Cat# ab14705, RRID:AB_2084810 |
| Mouse monoclonal anti-MT-CO2 | abcam | Cat# ab110258, RRID:AB_10887758 |
| Rabbit recombinant anti-Vinculin | abcam | Cat# ab219649, RRID:AB_2819348 |
| Rabbit monoclonal anti-ATF4 | Cell Signaling Technology | Cat# 11815, RRID:AB_2616025 |
| Rabbit monoclonal anti-CHOP | Cell Signaling Technology | Cat# 5554, RRID:AB_10694399 |
| Rabbit anti-cleaved caspase 3 | Cell Signaling Technology | Cat# 9661, RRID:AB_2341188 |
| Rabbit monoclonal anti-DELE1 | Cell Signaling Technology | Cat#98271 |
| Recombinant rabbit monoclonal anti-eIF2alpha | Cell Signaling Technology | Cat# 5324, RRID:AB_10692650 |
| Rabbit monoclonal anti-GAPDH (D16H11) | Cell Signaling Technology | Cat# 5174S, RRID: AB_10622025 |
| Rabbit monoclonal anti-LC3A/B | Cell Signaling Technology | Cat# 12741, RRID:AB_2617131 |
| Recombinant rabbit monoclonal anti-MFN2 | Cell Signaling Technology | Cat# 9482, RRID:AB_2716838 |
| Rabbit monoclonal anti-p-eIF2alpha | Cell Signaling Technology | Cat# 3398, RRID:AB_2096481 |
| Rabbit monoclonal anti-PINK1 | Cell Signaling Technology | Cat# 6946, RRID:AB_11179069 |
| Rabbit monoclonal anti-SOX2 | Cell Signaling Technology | Cat# 14962, RRID:AB_2798664 |
| Recombinant rabbit monoclonal anti-TOM20 | Cell Signaling Technology | Cat# 42406, RRID:AB_2687663 |
| HRP-linked anti-rabbit IgG | Cell Signaling Technology | Cat# 7074, RRID:AB_2099233 |
| HRP-linked anti-mouse IgG | Cell Signaling Technology | Cat# 7076, RRID:AB_330924 |
| Mouse monoclonal anti-OPA1 | BD Biosciences | Cat# 612606, RRID:AB_399888 |
| Mouse monoclonal anti-TOM20 | Santa Cruz Biotechnology | Cat# sc-17764, RRID:AB_628381 |
| Recombinant rabbit anti-HRI | Thermo Fisher Scientific | Cat# 711581, RRID:AB_2664567 |
| Rabbit polyclonal anti-GFAP | Proteintech | Cat# 16825-1-AP, RRID:AB_2109646 |
| Rabbit polyclonal anti-OLIG2 | Proteintech | Cat# 13999-1-AP, RRID:AB_2157541 |
| Rabbit polyclonal anti-p62/SQSTM1 | Proteintech | Cat# 18420-1-AP, RRID:AB_10694431 |
| Anti-mouse IgG Alexa Fluor 594 | Invitrogen | Cat# A11032 |
| Anti-rabbit IgG Alexa Fluor 594 | Invitrogen | Cat# A11012 |
| Anti-rabbit IgG Alexa Fluor 488 | Invitrogen | Cat# A11008 |
| **Chemicals** |  |  |
| Bafilomycin B1 | Cayman | 14005 |
| CellROX | Invitrogen | C10422 |
| Chloramphenicol | Thermo Fisher Scientific | 227920250 |
| Chloroquine diphosphate | Thermo Fisher Scientific | J64459.14 |
| Cyclosporin A | MedChemExpress | HYB0579 |
| Doxycycline hyclate | Sigma | D9891 |
| 2-deoxy-D-Glucose | Cayman | 14325 |
| FCCP | MedChemExpress | HY100410 |
| GCN2iB (GCN2 inhibitor) | MedChemExpress | HY112654 |
| GSK2656157 (PERK inhibitor) | MedChemExpress | HY13820 |
| Hemin | Sigma | H9039 |
| Hydrogen peroxide solution | Sigma | H1009 |
| Linezolid | LKT laboratories | 25912079 |
| MitoSOX | Invitrogen | M36008 |
| MitoTEMPO | Cayman | 16621 |
| N-Acetyl-L-cystein | Sigma | A9165 |
| Oligomycin A | MedChemExpress | HY16589 |
| PRK-IN-C16 (PKR inhibitor) | Selleckchem | S9668 |
| P110 | bio-technie TOCRIS | 6897 |
| TMRE | Thermo Fisher Scientific | T669 |
| Trans-ISRIB | bio-technie TOCRIS | 5284 |
| Rotenone | Sigma | R8875 |

**Table S2.**
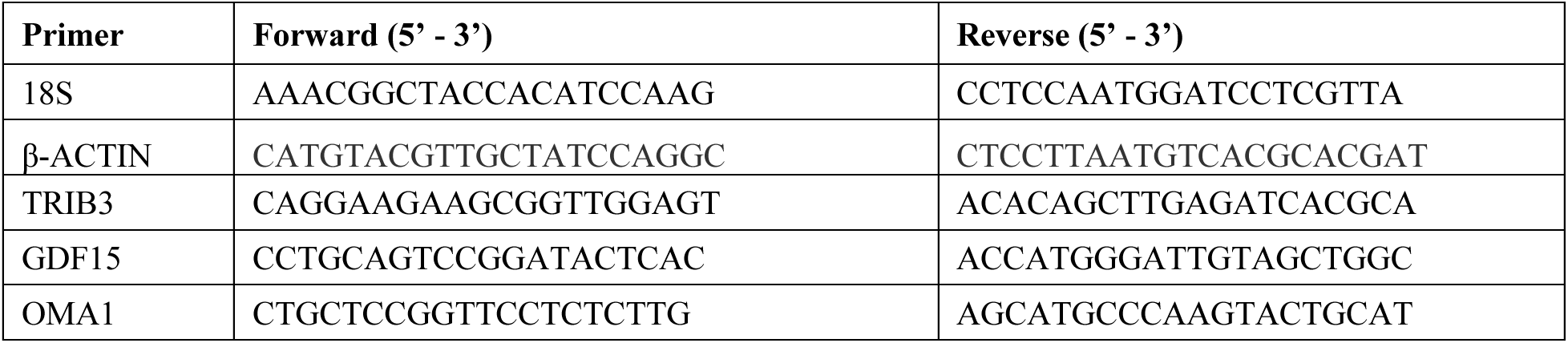
qPCR primer sequences.

